# PQS and Pyochelin in *Pseudomonas aeruginosa* Share Inner Membrane Transporters to Mediate Iron Uptake

**DOI:** 10.1101/2023.08.30.555523

**Authors:** Heng Zhang, Jianshe Yang, Juanli Cheng, Jing Zeng, Xin Ma, Jinshui Lin

## Abstract

Bacteria uptake different forms of iron through various channels to meet their needs. Our previous studies have shown that TseF, a type VI secretion system effector for Fe uptake, facilitates the delivery of outer membrane vesicle (OMV)-associated PQS-Fe^3+^ to bacterial cells by involving the Fe(III) pyochelin receptor FptA and the porin OprF. However, the form in which the PQS-Fe^3+^ complex enters the periplasm and how it is taken up into the cytoplasm remain unclear. Here, we first demonstrate that the PQS-Fe^3+^ complex enters the cell directly through FptA or OprF. Next, we find that inner membrane transporters such as FptX, PchHI, and FepBCDG are not only necessary for *P. aeruginosa* to uptake PQS-Fe^3+^ and PCH-Fe^3+^, but also necessary for the virulence of *P. aeruginosa* toward *Galleria mellonella* larvae. Furthermore, we suggest that the function of PQS-Fe^3+^ (but not PQS)-mediated quorum-sensing regulation is dependent on FptX, PchHI, and FepBCDG. Additionally, the findings indicate that, unlike FptX, both FepBCDG and PchHI play no role in the autoregulatory loop involving PchR, but further deletion of *fepBCDG* and *pchHI* can reverse the inactive PchR phenotype caused by *fptX* deletion and reactivate the expression of the PCH pathway genes under iron-limited conditions. Finally, this work detected the interaction between FptX, PchHI, and FepBCDG, indicating that a larger complex could be formed to mediate uptake of PQS-Fe^3+^ and PCH-Fe^3+^. These results pave the way for a better understanding of the PQS and PCH iron uptake pathway, and provide future directions to tackle *P. aeruginosa* infections.

**IMPORTANCE:** Iron is a key factor for *P. aeruginosa* to break through the host’s defense system and successfully infect. To acquire the necessary iron from the host, *P. aeruginosa* has evolved a number of strategies, the most common being the synthesis, secretion, and uptake of siderophores such as pyoverdine, pyochelin, and the quorum-sensing signaling molecule PQS. However, despite intensive studies of the siderophore uptake pathways of *P. aeruginosa*, our understanding of how siderophores transport iron across the inner membrane into the cytoplasm is still far from complete. Here, we reveal that PQS and pyochelin in *P. aeruginosa* share inner membrane transporters such as FptX, PchHI and FepBCDG to mediate iron uptake. Meanwhile, PQS and pyochelin-mediated signaling operates to a large extent via these inner membrane transporters. Our study revealed an interesting phenomenon of shared uptake pathways between PQS and pyochelin, which will lead us to reexamine the role of these two molecules in the iron uptake and virulence of *P. aeruginosa*.

## INTRODUCTION

Iron is a vital nutrient involved in a wide range of enzymatic functions and biological processes that is essential for bacterial growth and virulence(1). Iron-containing proteins exert a variety of vital functions, such as intermediary metabolism, cellular respiration, oxygen transport, transcription regulation, and DNA repair(2). However, in the host environment, iron is not readily available to bacteria due to the low solubility of iron and the activity of host iron-binding proteins (transferrin and lactoferrin)(3). Bacteria have evolved a number of strategies to acquire the necessary iron, including the secretion of siderophores. Siderophores are small compounds produced by bacteria under iron-limited conditions. The role of siderophores is to scavenge ferric iron in the bacterial environment and shuttle it back into the bacteria(4-8). Ferrisiderophore complexes are recognized at the cell surface of gram-negative bacteria by specific outer membrane transporters called TonB-dependent transporters (TBDTs). The biological function of TBDTs is to import siderophore–iron complexes from the extracellular medium into the periplasm(6). Once the ferrisiderophore enters the periplasm, it binds to siderophore–periplasmic binding protein (PBP) associated with the ATP-binding cassette (ABC) transporter. Then, the ferrisiderophore is imported into the cytoplasm via the interaction of the ferrisiderophore–PBP complex with the permease components of the ABC transporter(5, 9-11).

*Pseudomonas aeruginosa* is a ubiquitous opportunistic pathogen that can cause disease in immunocompromised hosts(4). Its infection is associated with a high incidence rate and mortality of many diseases, including pneumonia, chronic obstructive pulmonary disease (COPD), respiratory infections, and cystic fibrosis (CF)(12). Iron is necessary for *P. aeruginosa* to successfully infect the host, and for this purpose *P. aeruginosa* effectively competes for iron through a variety of independent mechanisms, as follows: (a) producing pyoverdine (PVD) and pyochelin (PCH), two siderophores that bind ferric iron with different affinities prior to being transferred into bacterial cells via the TBDTs (13, 14); (b) the uptake of xenosiderophores (1, 15-19); (c) ingesting heme molecules from host heme proteins via two heme uptake systems (Has and Phu)(20); (d) producing phenazine to reduce the external Fe^3+^ to Fe^2+^, which is transported into the cell through the Feo ferrous ion uptake system(21); and (e) using the quorum-sensing signal molecule 2-heptyl-3- hydroxy-4(1*H*)-quinolone (also known as *Pseudomonas* quinolone signal, PQS) to uptake free Fe^3+^ in the external environment(4). In *P. aeruginosa*, the *pqsABCDE* operon is responsible for the synthesis of the PQS precursor 2-heptyl-4- hydroxyquinoline (HHQ). In this process, FAD-dependent mono-oxygenase PqsH catalyzes the hydroxylation of HHQ to produce PQS(22). In recent years, with the continuous deepening of PQS research, it has been found that the function of PQS is not only reflected in quorum-sensing regulation but also in other aspects, such as mediating the formation of outer membrane vesicles(23-30), mediating iron uptake(4, 30-32), regulating host immunity(30, 33-35), mediating cytotoxicity(30, 36-39), and regulating population behavior(36, 40, 41). These studies suggest that the quorum sensing mediated by PQS in *P. aeruginosa* is not its primary function; instead, PQS plays a role similar to that of allelopathic substances in chemical ecology.

Recently, our research team made a major breakthrough in the research of the function of PQS in iron uptake(4). PQS produced in the cell is secreted to the extracellular space through an unknown pathway, and then integrated into the outer membrane to participate in the formation of outer membrane vesicles (OMVs)(42). Under low-iron conditions, the PQS in OMVs forms a PQS-Fe^3+^ complex with extracellular Fe^3+^. Then, TseF (a type-VI secretion system effector for Fe uptake) is secreted by H3-T6SS (a type-VI secretion system). TseF binds and tugs the PQS-Fe^3+^ complex to the PCH-Fe^3+^ receptor FptA and the outer membrane porin OprF. Thus, it may help PQS-Fe^3+^ to enter the periplasm of *P. aeruginosa*(4). However, how PQS-Fe^3+^ is further transported from the periplasm to the cytoplasm remains unknown. In addition, in the reported PCH-mediated iron uptake strategy, the outer membrane receptor FptA is responsible for the transport of PCH-Fe^3+^ into the periplasm, and then into the cell via the inner membrane transporter FptX(43). FptX is a known PCH-Fe^3+^ inner membrane transporter(43). However, its ability to transport PCH-Fe^3+^ into cells is only about 50%(44). Recent studies have shown that PchHI, an ABC family inner membrane transporter encoding the ATPase domain that is co-expressed with *fptX* in the biosynthetic gene cluster of PCH, is also involved in the siderophore-free iron uptake via PCH into the bacterial cytoplasm(5). However, despite intensive studies on the PCH pathway, the understanding of PCH-Fe^3+^ uptake is still far from complete.

Herein, we reveal that PCH-Fe^3+^ and PQS-Fe^3+^ are co-transported across the inner membrane through FptX, PchHI, and FepBCDG. Further investigation showed that FptX, PchHI, and FepBCDG were also involved in the virulence of *P. aeruginosa*. This study highlights the important roles of FptX, PchHI, and FepBCDG in the ability of *P. aeruginosa* to mediate iron uptake and virulence. These findings contributes to the further understanding of the molecular mechanism through which *P. aeruginosa* absorbs iron ions.

## RESULTS

### FptX, PchHI, and FepBCDG are involved in iron uptake via PQS

In the previously proposed iron uptake mechanism mediated by *P. aeruginosa* PQS, PQS-Fe^3+^ enters cells through the TseF-FptA/OprF pathway(4). However, this research group has discovered two possible manners in which PQS-Fe^3+^ enters cells through the TseF-FptA/OprF pathway. One is that PQS-Fe^3+^ first transmits iron to the receptors FptA or OprF, and then diffuses into cells in the form of PQS. The other is that PQS-Fe^3+^ directly enters cells through the receptors FptA or OprF. To further determine which of these two possible pathways plays a main role, this study compared the differences in the regulation of the lectin gene *lecA* and pyocyanin synthesis gene *phzA1*(*phzA1B1C1D1G1*) with the exogenous addition of PQS and PQS-Fe^3+^ in the PAΔ3Fe (a mutant defective in the pyoverdin biosynthetic pathway (Δ*pvdA*), ferrous iron transport (Δ*feoB*), PCH synthetase (Δ*pchE*), and PQS-Fe^3+^ biosynthetic (PqsA)) and PAΔ3FeΔ*pqsA*Δ*fptA*Δ*oprF*Δ*tseF* strains, in which transport (TseF-FptA/OprF) pathway was deleted in the background of strain PAΔ3FeΔ*pqsA*(4). The results are shown in Fig. S1. The exogenous addition of PQS and PQS-Fe^3+^ significantly induced the expression of *phzA1* and *lecA* in strain PAΔ3FeΔ*pqsA*, and adding PQS to the PAΔ3FeΔ*pqsA*Δ*fptA*Δ*oprF*Δ*tseF* strain also significantly induced the expression of *phzA1* and *lecA* in the trypticase soy broth (TSB) medium. However, adding PQS-Fe^3+^ did not effectively activate the expression of *phzA1* and *lecA* in the PAΔ3FeΔ*pqsA*Δ*fptA*Δ*oprF*Δ*tseF* strain in TSB medium. These results suggest that the TseF-FptA/OprF pathway does not affect the diffusion of PQS into the cell, but it is necessary for *P. aeruginosa* to uptake PQS-Fe^3+^ and allow PQS to function as a signaling molecule, indicating that PQS-Fe^3+^ enters the cell directly through the TseF-FptA/OprF pathway in the PAΔ3Fe strain.

Although the outer membrane receptor of PQS-Fe^3+^ is known, the mechanism of PQS-Fe^3+^ transmembrane transport has not yet been revealed. FptA is a known PCH-Fe^3+^ outer membrane receptor protein that mediates the uptake of PCH-Fe^3+^(43). Because PQS-Fe^3+^ and PCH-Fe^3+^ share the outer membrane receptor protein, we speculate that *P. aeruginosa* may also share the inner membrane transporter when ingesting PQS-Fe^3+^ and PCH-Fe^3+^. FptX is a known inner membrane transporter that plays an important role in PCH-Fe^3+^ uptake(43). Therefore, this study tested the effect of *fptX* mutation on the uptake of PQS-Fe^3+^ in *P. aeruginosa*. The results are shown in Fig. 1. The results showed that compared with PAΔ3Fe, further deleting *fptX* had an inhibitory effect on the uptake of PQS-Fe^3+^ by *P. aeruginosa*. However, this inhibitory effect was weaker than that in the negative control strain PAΔ3FeΔ*fptA*Δ*oprF* (Fig. 1B), indicating that there were other inner membrane transporters mediating the intracellular transport of PQS-Fe^3+^. Coincidentally, recent studies have found that the ABC family inner membrane transporter PchHI is also involved in the uptake of PCH-Fe^3+^ by *P. aeruginosa*(5). Therefore, we hypothesize that the inner membrane transporter PchHI is also involved in the uptake of PQS-Fe^3+^ by *P. aeruginosa*. The results of growth curve analysis confirmed this assumption (Fig. 1A–C). Compared with PAΔ3FeΔ*fptX*, strain PAΔ3FeΔ*fptXpchHI* had a lower uptake ability for PQS-Fe^3+^ in *P. aeruginosa* (Fig. 1B). The above results indicate that FptX and PchHI are jointly involved in the uptake of PQS-Fe^3+^ in *P. aeruginosa*.

**Fig. 1.**
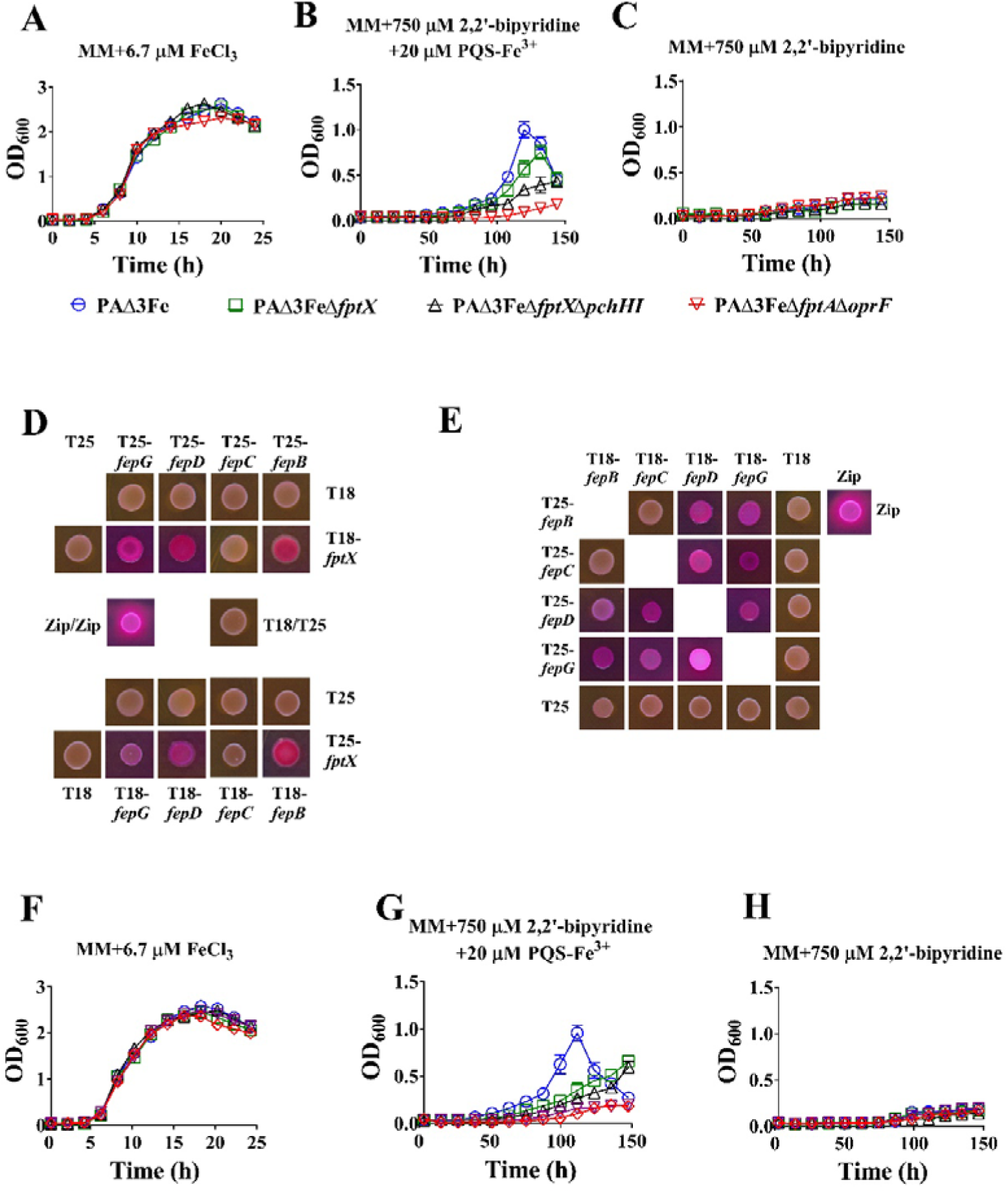
Effect of gene mutations *fptX*, *pchHI*, and *fepBCDG* on PQS-Fe^3+^ uptake by *Pseudomonas aeruginosa*. (A**–**C): Growth curves of *P. aeruginosa* mutant PAΔ3Fe and its derivative mutants (A) in the iron-limited succinate minimal medium (MM) supplemented with 6.7 μM FeCl_3_ (MM + 6.7μM FeCl_3_); (B) in the iron-limited MM supplemented with 750 μM iron chelator 2,2’- bipyridine and 20 μM PQS-Fe^3+^ (PQS:Fe^3+^ = 3:1) (MM + 750 μM 2,2’-bipyridine + 20 μM PQS-Fe^3+^); and (C) in the iron-limited MM supplemented with 750 μM iron chelator 2,2’-bipyridine (MM + 750 μM 2,2’-bipyridine). All the data represent the results of at least three independent experiments. The error bars represent the standard deviations. (D and E): Interactions between FptX, FepB, FepC, FepD, and FepG identified by bacterial two-hybrid experiments. Images of colonies formed by co-transformants on MacConkey agar plates (red colonies indicate a positive interaction). Various combinations of recombinant pKT25M and pUT18CM plasmids harboring proteins of interest were co-transformed into *Escherichia coli* BTH101, and the β-galactosidase activity of co-transformants was measured after plating on MacConkey agar plates (Fig. S2). Zip = leucine zipper domain of the yeast transcription factor GCN4 (positive control), T18 = empty vector pUT18CM, T25 = empty vector pKT25M. (F**–**H): Growth curves of *P. aeruginosa* PAΔ3Fe and its derivative mutants. The experimental conditions were similar to those in (A–C). All the data represent the results of at least three independent experiments. The error bars represent the standard deviations.

Although the ability of strain PAΔ3FeΔ*fptX*Δ*pchHI* to uptake PQS-Fe^3+^ was significantly reduced compared to PAΔ3Fe, it was still higher than that of the negative control strain PAΔ3FeΔ*fptA*Δ*oprF* (Fig. 1B), indicating that there were other inner membrane transporters mediating the uptake of PQS-Fe^3+^ in addition to FptX and PchHI. To identify other inner membrane transporters that may mediate the uptake of PQS-Fe^3+^, this study screened the proteins interacting with FptX from *P. aeruginosa* by constructing a bacterial two-hybrid screening library. The following four FptX-interacting proteins were identified: FepB (PA4159), FepC (PA4158), FepD (PA4160), and FepG (PA4161). These four proteins form the ABC family’s inner membrane transporter complex, FepBCDG(45). Bioinformatics predictions indicate that FepB is a periplasmic substrate-binding protein, FepC is an ATPase that carries an ATP-binding domain, and FepD and FepG are both cytoplasmic membrane permeases that may form heterodimers. Additionally, they may be involved in the uptake of enterobactin-Fe^3+^ in *P. aeruginosa*(7) (Table S3). To rule out false positives, this study further detected the interaction of full-length FepB, FepC, FepD, and FepG with FptX through bacterial two-hybrid assays. The results showed that FptX could interact with FepB, FepD and FepG, but could not interact with FepC (Figs. 1D and S2). In addition, this work further detected the interaction between FepB, FepC, FepD, and FepG. The results showed that there was no interaction between FepC and FepB, while other proteins could interact with each other (Figs. 1E and S2), indicating that FepB, FepC, FepD, and FepG together constituted the FepBCDG protein complex, which was consistent with the results of bioinformatics analysis. In conclusion, these results indicate that FptX interacts with the FepBCDG protein complex, which implies that the FepBCDG protein complex may also participate in the uptake of PQS-Fe^3+^ in *P. aeruginosa*.

To verify the above speculation, this work further deleted *fepBCDG* on the basis of PAΔ3FeΔ*fptX*Δ*pchHI* and detected its growth. The results showed that compared with strain PAΔ3FeΔ*fptX*Δ*pchHI*, the PAΔ3FeΔ*fptX*Δ*pchHI*Δ*fepBCDG* strain basically lost the ability to uptake PQS-Fe^3+^ to maintain normal growth. (Fig. 1F–H). This phenotype is similar to that of the negative control strain PAΔ3FeΔ*fptA*Δ*oprF*, indicating that the inner membrane transport process of *P. aeruginosa* to uptake PQS-Fe^3+^ is jointly mediated by FptX, PchHI, and FepBCDG. These results were also confirmed by genetic complementation (Fig. S3), which showed that the complementation of *fptX*, *pchHI*, and *fepBCDG* could promote the uptake of PQS-Fe^3+^ by PAΔ3FeΔ*fptX*Δ*pchHI*Δ*fepBCDG* to varying degrees.

### FptX, PchHI, and FepBCDG are involved in iron uptake via PCH

FptX is a known PCH-Fe^3+^ inner membrane transporter. However, it is only responsible for about 50% of the transport of PCH-Fe^3+^ into cells(44). In addition, the heterodimeric ABC transporter PchHI has also been reported to be involved in PCH-Fe^3+^ uptake by *P. aeruginosa*(5). Because FptX, PchHI, and FepBCDG are jointly involved in the uptake of PQS-Fe^3+^ in *P. aeruginosa*, they may also be jointly involved in the uptake of PCH-Fe^3+^. To verify this hypothesis, this study took the double deletion mutant Δ*pvdA*Δ*pqsH* as the starting strain, and the strain Δ*pvdA*Δ*pqsH*Δ*pchE*, with the simultaneous deletion of PVD, PQS, and PCH, as the negative control strain, and analyzed the effects of the deletion mutations *fptX*, *pchHI*, and *fepBCDG* on the uptake of PCH-Fe^3+^ in *P. aeruginosa*. The results are shown in Fig. 2A and B. Compared with Δ*pvdA*Δ*pqsH*, strain Δ*pvdA*Δ*pqsH*Δ*fptX* lost its partial ability to utilize PCH-Fe^3+^ to sustain normal growth. Knocking out the *pchHI* gene on the basis of the Δ*pvdA*Δ*pqsH*Δ*fptX* strain further reduced the ability of *P. aeruginosa* to uptake PCH-Fe^3+^. This result indicates that both FptX and PchHI play important roles in the uptake of PCH-Fe^3+^ in *P. aeruginosa.* However, the growth status of strain Δ*pvdA*Δ*pqsH*Δ*fptX*Δ*pchHI* was significantly better than that of the negative control strain Δ*pvdA*Δ*pqsH*Δ*pchE*. Therefore, on the basis of Δ*pvdA*Δ*pqsH*Δ*fptX*Δ*pchHI*, this study further deleted the *fepBCDG* gene. Compared with Δ*pvdA*Δ*pqsH*Δ*fptX*Δ*pchHI*, the growth phenotype of the Δ*pvdA*Δ*pqsH*Δ*fptX*Δ*pchHI*Δ*fepBCDG* strain was further inhibited. Consistent with the negative control strain Δ*pvdA*Δ*pqsH*Δ*pchE*, the strain Δ*pvdA*Δ*pqsH*Δ*fptX*Δ*pchHI*Δ*fepBCDG* completely lost the utilization of PCH-Fe^3+^. This finding suggests that FepBCDG also participates in the uptake of PCH-Fe^3+^ in *P. aeruginosa*. These results were also confirmed by the data on the results of genetic complementation (Fig. S4). In addition, the mutant Δ*pvdA*Δ*pqsH*Δ*pchE* was taken as the starting strain, and exogenous PCH or PCH extract (Δ*pvdA*Δ*pqsH* extract or Δ*pvdA*Δ*pqsH*Δ*pchE* extract) were added to the medium to analyze the effects of the deletion mutations *fptX*, *pchHI*, and *fepBCDG* on the uptake of PCH-Fe^3+^ in *P. aeruginosa*. The results are shown in Fig. 2C–G. Whether or not the Δ*pvdA*Δ*pqsH*Δ*pchE* extract was added to the iron-limited succinate minimal medium (MM), the growth of all strains was inhibited and was consistent with that of the negative control strain Δ*pvdA*Δ*pqsH*Δ*pchE*Δ*fptA*. However, the exogenous addition of Δ*pvdA*Δ*pqsH* extracts promoted the growth of all strains except the negative control strain Δ*pvdA*Δ*pqsH*Δ*pchE*Δ*fptA*, and among these strains, the positive control strain Δ*pvdA*Δ*pqsH*Δ*pchE* grew best (Fig. 2F). Compared with Δ*pvdA*Δ*pqsH*Δ*pchE*, Δ*pvdA*Δ*pqsH*Δ*pchE*Δ*fptX* lost its partial ability to sustain normal growth. Knocking out the *pchHI* gene on the basis of the Δ*pvdA*Δ*pqsH*Δ*pchE*Δ*fptX* strain further reduced the ability of *P. aeruginosa* to uptake PCH-Fe^3+^. However, compared with the negative control strain Δ*pvdA*Δ*pqsH*Δ*pchE*Δ*fptA*, growth was only partially affected in the strain Δ*pvdA*Δ*pqsH*Δ*pchE*Δ*fptX*Δ*pchHI*. The exogenous addition of PCH promoted the growth of all strains except for the negative control strain Δ*pvdA*Δ*pqsH*Δ*pchE*Δ*fptA* (Fig. 2G). Compared with Δ*pvdA*Δ*pqsH*Δ*pchE*, the Δ*pvdA*Δ*pqsH*Δ*pchE*Δ*fptX* strain lost its partial ability to utilize PCH-Fe^3+^ to sustain normal growth. Knocking out the *pchHI* gene on the basis of the Δ*pvdA*Δ*pqsH*Δ*pchE*Δ*fptX* strain further reduced the ability of *P. aeruginosa* to uptake PCH-Fe^3+^. However, compared with the negative control strain Δ*pvdA*Δ*pqsH*Δ*pchE*Δ*fptA*, growth was only partially affected in the strain Δ*pvdA*Δ*pqsH*Δ*pchE*Δ*fptX*Δ*pchHI*. Surprisingly, on the basis of Δ*pvdA*Δ*pqsH*Δ*pchE*Δ*fptX*Δ*pchHI*, after further deletion of the *fepBCDG* gene, strain Δ*pvdA*Δ*pqsH*Δ*pchE*Δ*fptX*Δ*pchHI*Δ*fepBCDG* was consistent with the negative control strain Δ*pvdA*Δ*pqsH*Δ*pchE*Δ*fptA*, and lost its growth promotion effect completely (Fig. 2F and G). It is suggested that FepBCDG is also involved in the uptake of PCH-Fe^3+^ by *P. aeruginosa*. These results were also confirmed by genetic complementation (Fig. S4). Overall, these data suggest that FptX, PchHI, and FepBCDG are jointly involved in the uptake of PCH-Fe^3+^ in *P. aeruginosa*.

**Fig. 2.**
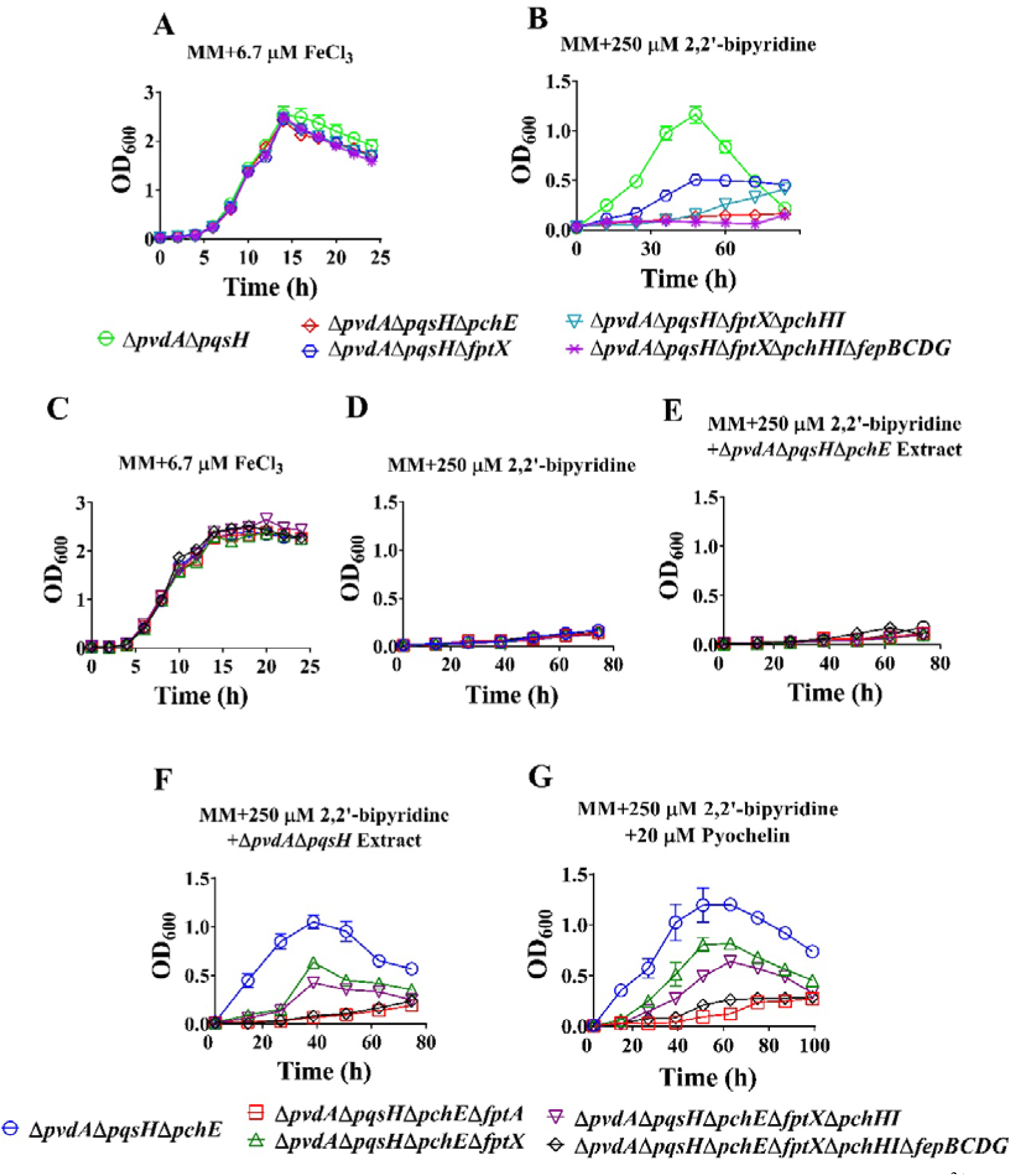
Effects of the deletion *fptX*, *pchHI*, and *fepBCDG* on the uptake of PCH-Fe^3+^ by *Pseudomonas aeruginosa*. (A, B): Growth curves of the *P. aeruginosa* mutant Δ*pvdA*Δ*pqsH* and its derivative mutants (A) in iron-limited succinate minimal medium (MM) supplemented with 6.7 μM FeCl_3_ (MM + 6.7 μM FeCl_3_), and (B) in iron-limited MM supplemented with 250 μM iron chelator 2,2’-bipyridine (MM+250 μM 2,2’-bipyridine). (C**–**G): Growth curves of *P. aeruginosa* Δ*pvdA*Δ*pqsH*Δ*pchE* and its derivative mutants in the iron-limited MM. (C) Experimental conditions were similar to those in A. (D) Experimental conditions were similar to those in B. (E) Addition of 250 μM 2,2’-bipyridine and Δ*pvdA*Δ*pqsH*Δ*pchE* extract. (F) Addition of 250 μM 2,2’-bipyridine and Δ*pvdA*Δ*pqsH* extract. (G) Addition of 250 μM 2,2’-bipyridine and 20 μM PCH. All the data represent the results of at least three independent experiments. The error bars represent the standard deviations.

### FptX, PchHI, and FepBCDG are necessary for PQS-Fe^3+^ and PCH-Fe^3+^ uptake

To analyze the roles of FptX, PchHI, and FepBCDG in the uptake of PQS-Fe^3+^ and PCH-Fe^3+^ in *P. aeruginosa*, PAΔ3FeΔ*pqsH* (or Δ*pvdA*Δ*pqsH*Δ*pchE*) was used as the starting strain to exclude the interference of *P. aeruginosa* from utilizing endogenous PVD, PCH, ferrous iron, and PQS (or PVD, PCH, and PQS) on the experimental results, and the growth differences between PAΔ3FeΔ*pqsH* and PAΔ3FeΔ*pqsH*Δ*fptX*Δ*pchHI*Δ*fepBCDG* or Δ*pvdA*Δ*pqsH*Δ*pchE* and Δ*pvdA*Δ*pqsH*Δ*pchE*Δ*fptX*Δ*pchHI*Δ*fepBCDG* in co-culture were compared. The results showed that the growth ability of PAΔ3FeΔ*pqsH*Δ*fptX*Δ*pchHI*Δ*fepBCDG* or Δ*pvdA*Δ*pqsH*Δ*pchE*Δ*fptX*Δ*pchHI*Δ*fepBCDG* in the iron-limited MM supplemented with PQS-Fe^3+^ or PCH was significantly lower than that of PAΔ3FeΔ*pqsH* or Δ*pvdA*Δ*pqsH*Δ*pchE* in the same conditions. However, in the iron-limited MM with the addition of hemin as a control, the growth ability of PAΔ3FeΔ*pqsH* and PAΔ3FeΔ*pqsH*Δ*fptX*Δ*pchHI*Δ*fepBCDG* or Δ*pvdA*Δ*pqsH*Δ*pchE* and Δ*pvdA*Δ*pqsH*Δ*pchE*Δ*fptX*Δ*pchHI*Δ*fepBCDG* was consistent (Fig. 3A–D), suggesting that *fptX*, *pchHI*, and *fepBCDG* mutations significantly reduced the ability of PAΔ3FeΔ*pqsH* and Δ*pvdA*Δ*pqsH*Δ*pchE* to utilize PQS-Fe^3+^ and PCH, but did not affect their ability to utilize hemin under iron-limiting growth conditions. These results indicate that the *fptX*, *pchHI*, and *fepBCDG* genes are crucial for *P. aeruginosa* to utilize PQS-Fe^3+^ or PCH-Fe^3+^.

**Fig. 3.**
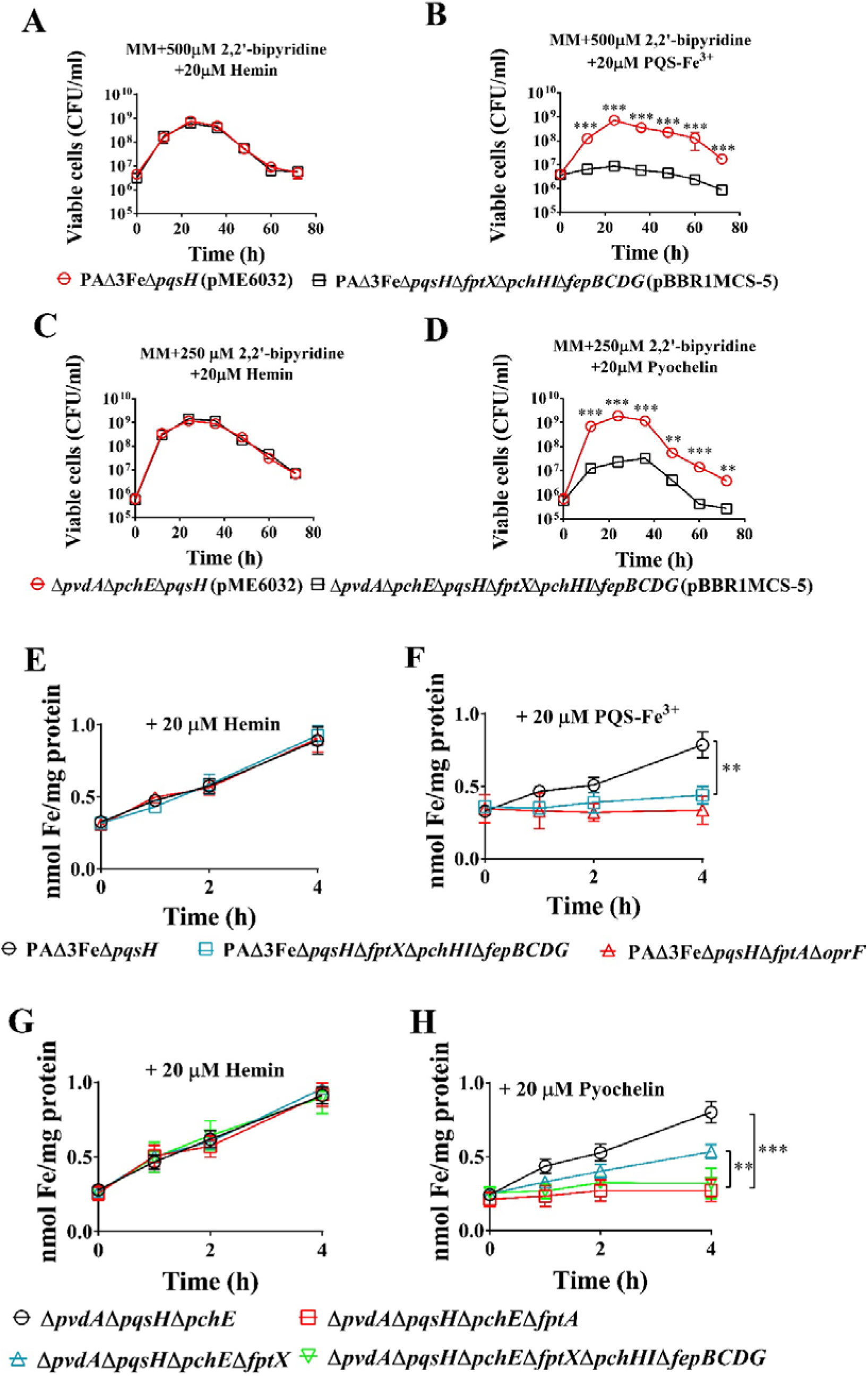
Effects of the deletion of *fptX*, *pchHI*, and *fepBCDG* on the utilization of PQS-Fe^3+^ and PCH-Fe^3+^ by *Pseudomonas aeruginosa*. (A, B): Co-culture growth curves of *P. aeruginosa* mutants PAΔ3FeΔ*pqsH* and PAΔ3FeΔ*pqsH*Δ*fptX*Δ*pchHI*Δ*fepBCDG* in iron-limited succinate minimal medium (MM). (A) Addition of 500 μM 2,2’-bipyridine and 20 μM hemin to the iron-limited MM. (B) Addition of 500 μM 2,2’-bipyridine and 20 μM PQS-Fe^3+^ (PQS:Fe^3+^ = 3:1) to the iron-limited MM. (C, D) Co-culture growth curves of *P. aeruginosa* mutants Δ*pvdA*Δ*pqsH*Δ*pchE* and Δ*pvdA*Δ*pqsH*Δ*pchE*Δ*fptX*Δ*pchHI*Δ*fepBCDG* in iron-limited MM. (C) Addition of 250 μM 2,2’-bipyridine and 20 μM hemin to the iron-limited MM. (D) Addition of 250 μM 2,2’-bipyridine and 20 μM pyochelin to the iron-limited MM. (E, F) *P. aeruginosa* mutants PAΔ3FeΔ*pqsH* and PAΔ3FeΔ*pqsH*Δ*fptX*Δ*pchHI*Δ*fepBCDG* were cultured in iron-limited MM to the mid-log phase. Cells were collected and suspended in phosphate-buffered saline (PBS), 0.4% glucose and 20 µM hemin or PQS-Fe^3+^ (PQS:Fe^3+^ = 3:1) were added, and the solutions were incubated at 37°C and 200 rpm for 0 h, 1 h, 2 h, and 4 h. The cell samples were collected and the intracellular metal ion content was determined using inductively coupled plasma mass spectrometry (ICP-MS). (E) Addition of 20 µM hemin to PBS. (F) Addition of 20 μM PQS-Fe^3+^ (PQS:Fe^3+^=3:1) to PBS. The PAΔ3FeΔ*pqsH*Δ*fptA*Δ*oprF* strain served as the negative control. (G, H) *P. aeruginosa* mutants Δ*pvdA*Δ*pqsH*Δ*pchE*, Δ*pvdA*Δ*pqsH*Δ*pchE*Δ*fptX*, and Δ*pvdA*Δ*pqsH*Δ*pchE*Δ*fptX*Δ*pchHI*Δ*fepBCDG* were cultured in iron-limited MM to the mid-log phase. Cells were collected and suspended in PBS, 0.4% glucose and 20 µM hemin or pyochelin were added, and the solution was incubated at 37°C and 200 rpm for 0 h, 1 h, 2 h, and 4 h. The cell samples were collected and the intracellular metal ion content was determined using ICP-MS. (G) Experimental conditions were similar to those in E. (H) Addition of 20 μM PCH to PBS. The Δ*pvdA*Δ*pqsH*Δ*pchE*Δ*fptA* strain served as the negative control. All data represent the results of at least three independent experiments. Error bars represent standard deviations. **, *P* < 0.01, ***, *P* < 0.001.

To further analyze the roles of FptX, PchHI, and FepBCDG in the uptake of PQS-Fe^3+^ and PCH-Fe^3+^, this study firstly cultured strains PAΔ3FeΔ*pqsH* and PAΔ3FeΔ*pqsH*Δ*fptX*Δ*pchHI*Δ*fepBCDG* (for the analysis of PQS-Fe^3+^ uptake), or strains Δ*pvdA*Δ*pqsH*Δ*pchE*, Δ*pvdA*Δ*pqsH*Δ*pchE*Δ*fptX*, and Δ*pvdA*Δ*pqsH*Δ*pchE*Δ*fptX*Δ*pchHI*Δ*fepBCDG* (for the analysis of PCH-Fe^3+^ uptake) in the iron-limited MM to the mid-log phase, and then subcultured these strains in phosphate-buffered saline (PBS) (containing 0.4% glucose) supplemented with PQS-Fe^3+^, PCH, or hemin. Cell samples were collected at different time points to measure the content of intracellular metal ions. The results are shown in Fig. 3E–H. Under the treatment conditions of the exogenous addition of PQS-Fe^3+^ or PCH, the intracellular iron content of strains PAΔ3FeΔ*pqsH* or Δ*pvdA*Δ*pqsH*Δ*pchE* increased rapidly with the extension of culture time; meanwhile, the intracellular iron content of strains PAΔ3FeΔ*pqsH*Δ*fptX*Δ*pchHI*Δ*fepBCDG* and Δ*pvdA*Δ*pqsH*Δ*pchE*Δ*fptX*Δ*pchHI*Δ*fepBCDG* changed little and was consistent with that of the negative control strain PAΔ3FeΔ*pqsH*Δ*fptA*Δ*oprF* and Δ*pvdA*Δ*pqsH*Δ*pchE*Δ*fptA*, and significantly lower than that of strains PAΔ3FeΔ*pqsH* and Δ*pvdA*Δ*pqsH*Δ*pchE* (Fig. 3F and H). However, under the same conditions, there were no significant differences in the intracellular zinc and manganese ion contents in these strains (Fig. S5). Additionally, under the treatment condition of the exogenous addition of hemin, the intracellular iron, zinc, and manganese ion contents exhibited no significant differences among these strains (Figs. 3E, 3G, and S5). In addition, complementary *fptX*, *pchHI*, or *fepBCDG* genes could promote the increase of intracellular iron content in the corresponding mutant cells to varying degrees under the treatment conditions of the exogenous addition of PQS-Fe^3+^ or PCH (Fig. S5). However, complementary *fptX*, *pchHI*, or *fepBCDG* genes had no effect on the contents of intracellular zinc and manganese ions under the same conditions (Fig. S5). These results suggested that the mutation of *fptX*, *pchHI*, and *fepBCDG* significantly reduced the uptake of PQS-Fe^3+^ and PCH-Fe^3+^ in the PAΔ3FeΔ*pqsH* and Δ*pvdA*Δ*pqsH*Δ*pchE* strains, respectively. In summary, FptX, PchHI, and FepBCDG are necessary for *P. aeruginosa* to uptake PQS-Fe^3+^ and PCH-Fe^3+^.

### Lack of an energy source impairs the uptake of PQS-Fe^3+^ and PCH-Fe^3+^ by *P. aeruginosa* inner membrane transporters

We have demonstrated that FptX, PchHI, and FepBCDG are involved in the uptake of PQS-Fe^3+^ or PCH-Fe^3+^. Interestingly, FptX is a proton motive-dependent permease(7, 44). The PchH and PchI proteins carry an ATP-binding domain and a transmembrane domain, respectively, on the same polypeptide(46, 47), and the two proteins may function together to form a complete heterodimeric ABC transporter(5, 46). Therefore, this study analyzed the effects of lacking an energy source on the functions of PQS-Fe^3+^ and PCH-Fe^3+^ inner membrane transporters. For FptX, which uses proton motive force as its energy source, carbonyl cyanide-m-chlorophenylhydrazone (CCCP) was used to inhibit the cell’s proton motive force and cause it to lose its energy source. Under the iron-limited culture conditions of adding PQS-Fe^3+^, the growth ability of PAΔ3FeΔ*pchHI*Δ*fepBCDG* was close to that of PAΔ3Fe, and was significantly higher than that of PAΔ3FeΔ*fptX*Δ*pchHI*Δ*fepBCDG*. However, when 10 µM CCCP was added to the medium, the growth of both the PAΔ3FeΔ*pchHI*Δ*fepBCDG* and PAΔ3FeΔ*fptX*Δ*pchHI*Δ*fepBCDG* strains was completely inhibited, while PAΔ3Fe still grew effectively (Fig. 4A). In addition, in the Δ*pvdA*Δ*pqsH* mutant and its derivative strains (for analyzing the role of endogenous PCH), the growth ability of Δ*pvdA*Δ*pqsH*Δ*pchHI*Δ*fepBCDG* was close to that of Δ*pvdA*Δ*pqsH*, and was significantly higher than that of Δ*pvdA*Δ*pqsH*Δ*fptX*Δ*pchHI*Δ*fepBCDG* in the iron-limited MM. However, when 250 µM CCCP was added to the medium, the growth of both the Δ*pvdA*Δ*pqsH*Δ*pchHI*Δ*fepBCDG* and Δ*pvdA*Δ*pqsH*Δ*fptX*Δ*pchHI*Δ*fepBCDG* strains was completely inhibited, while Δ*pvdA*Δ*pqsH* still grew effectively (Fig. 4B), indicating that CCCP inhibited the energy source of FptX, resulting in its loss of the ability to transport PQS-Fe^3+^ and PCH-Fe^3+^. For PchH and PchI, which use ATP as an energy source, the conserved amino acid residues on the Walker B motif of their ATPase domain exhibited site-directed mutation, and their mutation sites were both E490A, so that they lost ATPase activity and their energy source (Fig. 4C). As shown in Fig. 4D, compared with the complementary wild-type gene *pchHI*, complementary *pchH*\**I* (i.e. PchH^E490A^I) could not restore the growth of PAΔ3FeΔ*fptX*Δ*pchHI*Δ*fepBCDG* in the iron-limited MM. However, the growth phenotype of the PAΔ3FeΔ*fptX*Δ*pchHI*Δ*fepBCDG* strains with complementary *pchHI\** (i.e. PchHI^E490A^) and complementary wild-type *pchHI* were similar. In addition, a similar situation was observed in the genetic complementary strains of strain Δ*pvdA*Δ*pqsH*Δ*fptX*Δ*pchHI*Δ*fepBCDG*. Compared with the complementary wild-type *pchHI*, complementary *pchH*\**I* (i.e. PchH^E490A^I) could not restore the growth of Δ*pvdA*Δ*pqsH*Δ*fptX*Δ*pchHI*Δ*fepBCDG* in the iron-limited MM. However, the growth phenotypes of Δ*pvdA*Δ*pqsH*Δ*fptX*Δ*pchHI*Δ*fepBCDG* strains with complementary *pchHI\** (i.e. PchHI^E490A^) and complementary wild-type *pchHI* were similar (Fig. 4E). These results indicated that the energy of the PchHI transporter complex was derived from the ATPase activity of PchH, but not PchI. FepC provides energy for the ABC type transporter complex FepBCDG. The energy source of FepC was lost through the site-directed mutagenesis of the conserved amino acid residue on the Walker B motif of the FepC ATPase domain (the mutation site was E166A) (Fig. 4C). The results are shown in Fig. 4F and G. Compared with the complementary wild type gene *fepC*, the complementary *fepC*\* (i.e. FepC^E166A^) could not restore the growth of the PAΔ3FeΔ*fptX*Δ*pchHI*Δ*fepC* strain in the iron-limited MM, suggesting that the energy of the FepBCDG transporter complex was derived from the ATPase activity of FepC. To sum up, these results indicate that the lack of an energy source impairs the capacity of the *P. aeruginosa* inner membrane transporters to transport PQS-Fe^3+^ and PCH-Fe^3+^.

**Fig. 4.**
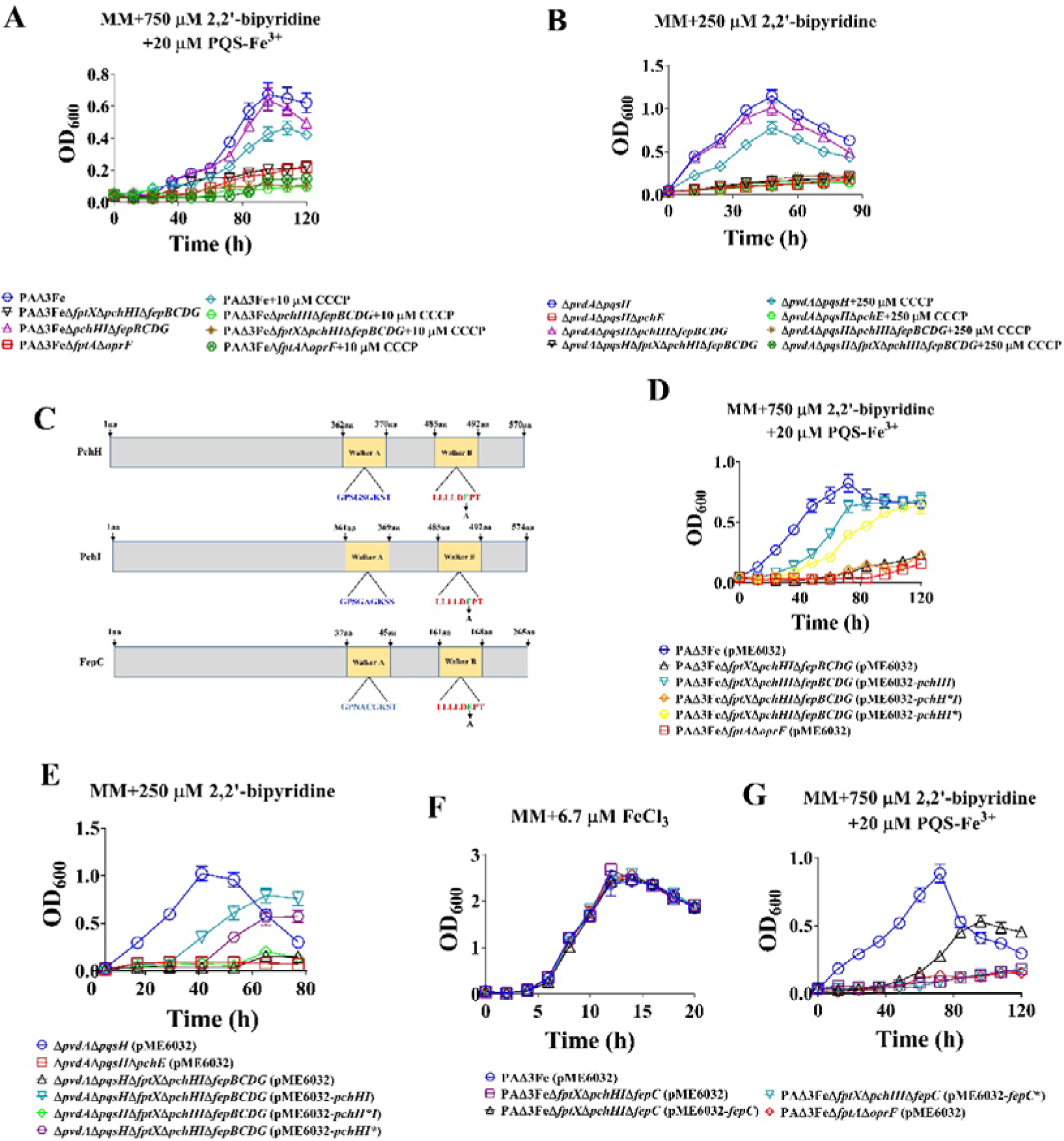
Effects of energy source deficiency on the uptake of PQS-Fe^3+^ and PCH-Fe^3+^ in *Pseudomonas aeruginosa.* (A) Growth curves of *P. aeruginosa* PAΔ3Fe and its derivative mutants in iron-limited succinate minimal medium (MM) supplemented with 750 μM 2,2’-bipyridine and 20 μM PQS-Fe^3+^ (PQS:Fe^3+^ = 3:1) and the presence or absence of 10 μM carbonyl cyanide-m-chlorophenylhydrazone (CCCP). (B) Growth curves of *P. aeruginosa* Δ*pvdA*Δ*pqsH* and its derivative mutants in iron-limited MM supplemented with 250 μM 2,2’-bipyridine and the presence or absence of 250 μM CCCP. (C) Schematic diagram of the PchH, PchI, and FepC ATPase domains. (D) Growth curve of complementation strains of *P. aeruginosa* PAΔ3FeΔ*fptX*Δ*pchHI*Δ*fepBCDG* in iron-limited MM supplemented with 750 μM 2,2’-bipyridine and 20 μM PQS-Fe^3+^ (PQS:Fe^3+^ = 3:1). (E) Growth curve of complementation strains of *P. aeruginosa* PAΔ3FeΔ*fptX*Δ*pchHI*Δ*fepBCDG* in iron-limited MM supplemented with 250 μM 2,2’-bipyridine. (F, G) Growth curve of complementation strains of *P. aeruginosa* PAΔ3FeΔ*fptX*Δ*pchHI*Δ*fepC* in iron-limited MM. (F) Addition of 6.7 μM FeCl_3_ to the iron-limited MM. (G) Addition of 750 μM 2,2’-bipyridine and 20 μM PQS-Fe^3+^ to the iron-limited MM. All data represent the results of at least three independent experiments. Error bars represent standard deviations.

### FptX, PchHI, and FepBCDG affect the expression of the lectin gene *lecA* and the PCH biosynthetic operon

The PQS signaling molecule in *P. aeruginosa* regulates the expression of many virulence genes by combining the transcription regulator PqsR, including the *pqsABCDE* synthesis operon, the pyocyanin *phzA1B1C1D1G1* synthesis operon, and the lectin *lecA* gene(22). In *P. aeruginosa* PQS-deficient mutant Δ*pqsA*, the exogenous addition of PQS significantly induced the expression of *pqsA* and *lecA* gene in iron-sufficient Luria–Bertani (LB) medium, while the exogenous addition of PQS or PQS-Fe^3+^ (PQS: Fe^3+^ = 3:1) could also effectively induce the expression of the *pqsA* gene in iron-deficient casamino acid (CAA) medium, indicating that the iron-chelating activity of PQS did not affect the function of PQS signal molecules under this specific condition(38). Because FptX, PchHI, and FepBCDG, three inner membrane transporters, participate in the uptake of PQS-Fe^3+^ in *P. aeruginosa*, do they also affect the function of PQS as a quorum-sensing signal molecule? Using the *lecA* gene promoter, which is regulated by PQS, as a probe, this study compared the differences in the regulation of the lectin gene *lecA* by the exogenous addition of PQS and PQS-Fe^3+^ in the PAΔ3FeΔ*pqsA* and PAΔ3FeΔ*pqsA*Δ*fptX*Δ*pchHI*Δ*fepBCDG* strains. The results are shown in Fig. 5A. The exogenous addition of PQS did not significantly activate *lecA* in the iron-limited medium (the TSB medium containing 300 μM 2,2’-bipyridine), but significantly activated *lecA* in the TSB medium (Fig. S1), indicating that under iron-limited conditions, PQS was not effective for uptake by *P. aeruginosa*. On the contrary, when PQS-Fe^3+^ was added, the expression of the *lecA* gene was only significantly activated in the PAΔ3FeΔ*pqsA* strain, while it was not activated in the PAΔ3FeΔ*pqsA*Δ*fptX*Δ*pchHI*Δ*fepBCDG* strain, indicating that under iron-limited culture conditions, PQS mainly entered cells in the form of PQS-Fe^3+^, and this process depended on FptX, PchHI, and FepBCDG.

**Fig. 5.**
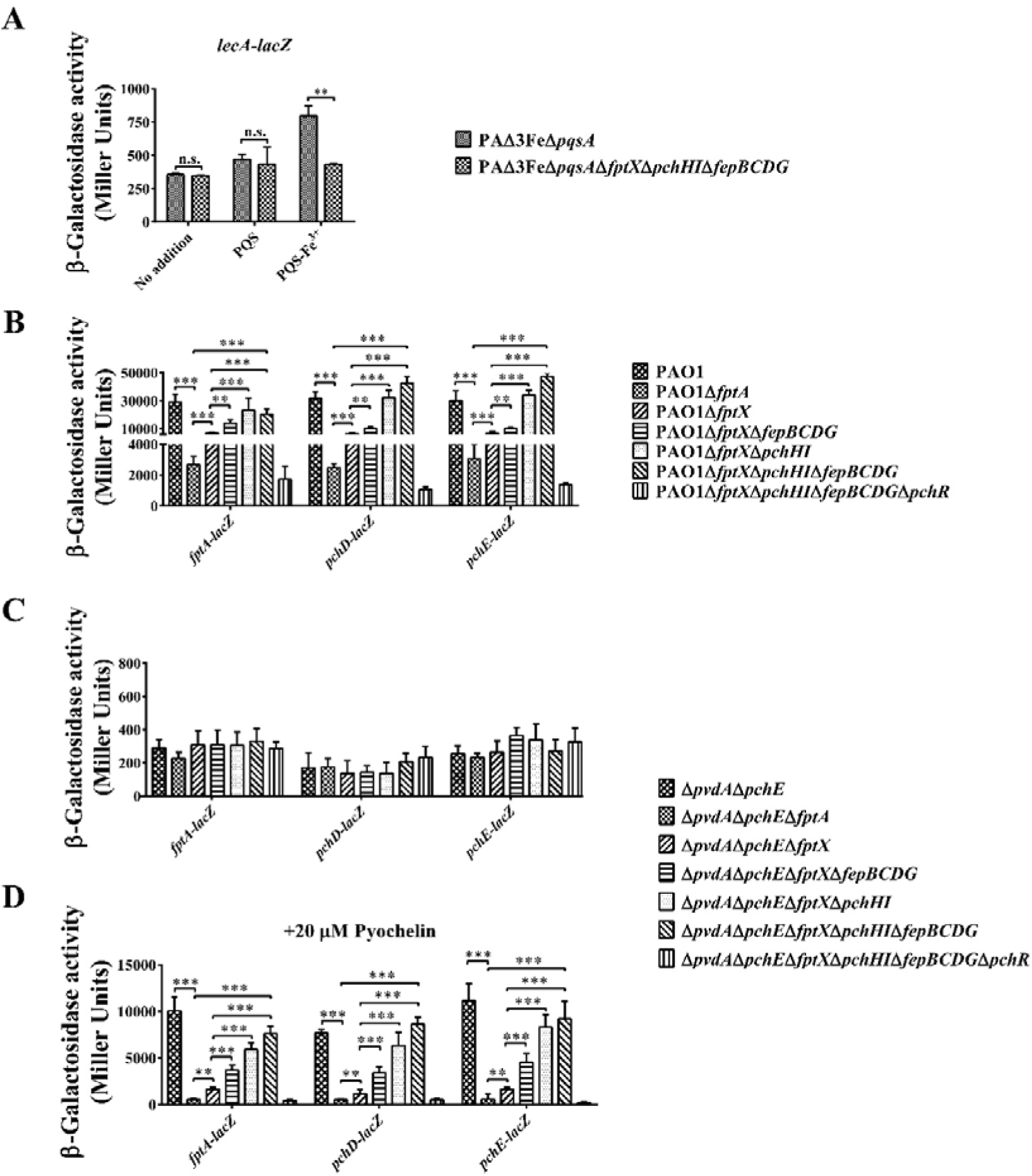
Effects of the deletion of *fptX*, *pchHI*, and *fepBCDG* on the expression of the lectin *lecA* gene and PCH genes in *Pseudomonas aeruginosa*. (A) Cells cultured in iron-limited medium (trypticase soy broth (TSB) medium containing 300 μM 2,2’-bipyridine) were supplied with or without 40 μM PQS or PQS-Fe^3+^ (PQS:Fe^3+^ = 3:1), and the transcription levels of *lecA* in *P. aeruginosa* PAΔ3FeΔ*pqsA* and PAΔ3FeΔ*pqsA*Δ*fptX*Δ*pchHI*Δ*fepBCDG* mutant cells were monitored using the *lecA–lacZ* transcriptional fusions. (B) Cells were cultured in iron-limited succinate minimal medium (MM) and the levels of *fptABCX*, *pchDCBA*, and *pchEFGHI* transcription in *P. aeruginosa* PAO1, PAO1Δ*fptA*, PAO1Δ*fptX*, PAO1Δ*fptX*Δ*fepBCDG*, PAO1Δ*fptX*Δ*pchHI*, PAO1Δ*fptX*Δ*pchHI*Δ*fepBCDG*, and PAO1Δ*fptX*Δ*pchHI*Δ*fepBCDG*Δ*pchR* mutant cells were monitored using the *fptA–lacZ*, *pchD–lacZ*, and *pchE–lacZ* transcriptional fusions, respectively. (C and D) Cells cultured in iron-limited MM were supplied with or without 20 μM pyochelin and the transcription levels of *fptABCX*, *pchDCBA*, and *pchEFGHI* in *P. aeruginosa* Δ*pvdApchE*, Δ*pvdApchE*Δ*fptA*, Δ*pvdApchE*Δ*fptX*, Δ*pvdApchE*Δ*fptX*Δ*fepBCDG*, Δ*pvdApchE*Δ*fptX*Δ*pchHI*, Δ*pvdApchE*Δ*fptX*Δ*pchHI*Δ*fepBCDG*, and Δ*pvdApchE*Δ*fptX*Δ*pchHI*Δ*fepBCDG*Δ*pchR* mutant cells were monitored using the *fptA–lacZ*, *pchD–lacZ*, and *pchE–lacZ* transcriptional fusions, respectively. (C) No addition of pyochelin to the iron-limited MM. (D) Addition of 20 μM pyochelin to the iron-limited MM. The graphs show the mean and standard deviation of three experiments performed in five replicates each time. n.s., not significant, **, *P* < 0.01, ***, *P* < 0.001.

The siderophore PCH of *P. aeruginosa* regulates the expression of many genes via combining the transcription regulator PchR, including the PCH biosynthesis operons *pchDCBA* and *pchEFGHI*, and the PCH-Fe^3+^ uptake operon *fptABCX*(44, 48). Because FptX, PchHI, and FepBCDG, three inner membrane transporters, participate in the uptake of PCH-Fe^3+^ in *P. aeruginosa*, and both FptA and FptX are involved in the positive autoregulatory loop through importing the PCH-Fe^3+^ complex interacting with PchR into the bacteria(5, 44), do PchHI and FepBCDG also play important roles in PCH-mediated signaling? To answer this question, we used the *fptA*, *pchD*, and *pchE* gene promoters as probes to analyze the role of PCH-mediated signaling by FptX, PchHI, and FepBCDG through *lacZ* transcription fusion of the promoter. *P. aeruginosa* PAO1 was used as the starting strain to analyze the regulation of these genes by endogenous PCH. Consistent with previous reports(44), when the *fptA* or *fptX* gene was deleted, the expression of *fptA*, *pchD*, and *pchE* was significantly down-regulated under iron-limited conditions. However, when further deletion of *fepBCDG* was performed on the basis of PAO1Δ*fptX*, the expression of the *fptA*, *pchD*, and *pchE* genes was not affected. In contrast to the single PAO1Δ*fptX* mutant, the PAO1Δ*fptX*Δ*pchHI* and PAO1Δ*fptX*Δ*pchHI*Δ*fepBCDG* mutants showed the phenotype to activate expression of the PCH genes. As expected, the deletion of *pqsR* on the basis of PAO1Δ*fptX*Δ*pchHI*Δ*fepBCDG* had a strong inhibitory effect on the expression of the genes of the PCH locus, indicating that the positive autoregulatory loop involving PchR was no longer active (Fig. 5B). In addition, Δ*pvdA*Δ*pchE* and its derivative mutants were used to study how exogenous PCH regulated the expression of the PCH genes. The results showed that when no exogenous PCH was added to the medium, there were only background gene expression levels and no significant differences in the expression of the *fptA*, *pchD*, and *pchE* genes between Δ*pvdA*Δ*pchE* and its derivative mutants (Fig. 5C). In contrast, when PCH was added, the expression levels of the *fptA*, *pchD*, and *pchE* genes were significantly up-regulated in the Δ*pvdA*Δ*pchE* mutant. As expected, after the deletion of the *fptA* or *fptX* genes on the basis of the Δ*pvdA*Δ*pchE* mutant, the exogenous addition of PCH failed to activate the expression of *fptA*, *pchD*, and *pchE*. Similarly, the deletion of *pqsR* on the basis of the Δ*pvdA*Δ*pchE*Δ*fptX*Δ*pchHI*Δ*fepBCDG* mutant made the expression of the PCH genes no longer responsive to exogenously added PCH. However, in contrast to the Δ*pvdA*Δ*pchE*Δ*fptX* mutant, the Δ*pvdA*Δ*pchE*Δ*fptX*Δ*fepBCDG*, Δ*pvdA*Δ*pchE*Δ*fptX*Δ*pchHI*, and Δ*pvdA*Δ*pchE*Δ*fptX*Δ*pchHI*Δ*fepBCDG* mutants exhibited the phenotype to activate the expression of the PCH genes under the same culture conditions (Fig. 5D). These results demonstrate that, unlike FptX, both FepBCDG and PchHI play no role in the autoregulatory loop involving PchR, but further deletion of *fepBCDG* and *pchHI* can reverse the inactive PchR phenotype caused by *fptX* deletion and reactivate the expression of the genes of the PCH pathway under iron-limited conditions.

### FptX, PchHI, and FepBCDG are necessary for the virulence of *P. aeruginosa* in *Galleria mellonella* larvae

PQS and PCH play important roles in the virulence of *P. aeruginosa*(49, 50), and FptX, PchHI, and FepBCDG, three inner membrane transporters, jointly participate in the uptake of PQS-Fe^3+^ and PCH-Fe^3+^ in *P. aeruginosa*. Thus, we speculated that FptX, PchHI, and FepBCDG may affect the virulence of *P. aeruginosa*. To test this hypothesis, this study analyzed the effect of these inner membrane transporter deletions on the toxicity of *P. aeruginosa* to *G. mellonella* larvae and the viability of *P. aeruginosa* in the *G. mellonella* larvae. *G. mellonella* is a host that can be used to examine the virulence of this pathogen(4). The Δ*pvdA* mutant strain was used as the starting strain to eliminate the interference of siderophore PVD. The results showed that the toxicity of strain Δ*pvdA*Δ*fptX*Δ*pchHI*Δ*fepBCDG* to the *G. mellonella* larvae was significantly lower than that of the Δ*pvdA* strain (Fig. 6A), while the complementary *fptX*, *pchHI*, and *fepBCDG* recovered the toxicity of *P. aeruginosa* to varying degrees (Fig. S6). This study further evaluated the role of PQS-mediated or PCH-mediated iron uptake in the biology of *P. aeruginosa* by examining the interactions of relevant mutants with *G. mellonella*. Mutants defective in *fptX*, *pchHI*, and *fepBCDG* were able to effectively compete with the parental strain PAΔ3Fe or Δ*pvdA*Δ*pqsH* in media. The deletion of *fptX*, *pchHI*, or *fepBCDG* from mutant PAΔ3Fe or Δ*pvdA*Δ*pqsH* resulted in mutants that could not effectively compete against PAΔ3Fe or Δ*pvdA*Δ*pqsH* when co-inoculated in the host (Fig. 6B and C). However, complementing *fptX*, *pchHI*, and *fepBCDG* could restore the survival competition-deficient phenotype of PAΔ3FeΔ*fptX*Δ*pchHI*Δ*fepBCDG* or Δ*pvdA*Δ*pqsH*Δ*fptX*Δ*pchHI*Δ*fepBCDG* in the *G. mellonella* larvae to varying degrees (Fig. S6). These results indicate that FptX, PchHI, and FepBCDG are necessary for *P. aeruginosa* virulence in this host.

**Fig. 6.**
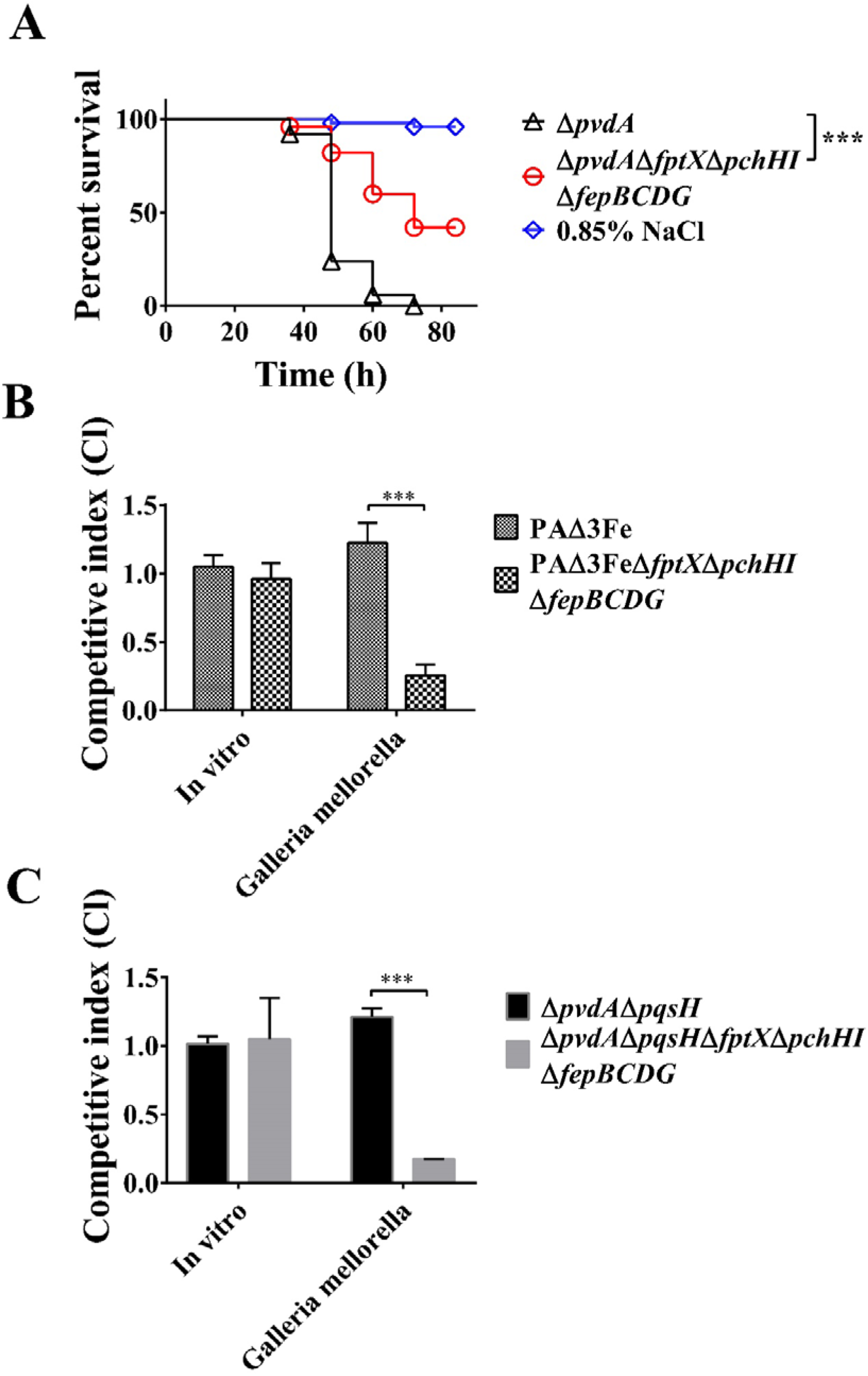
Deletion of *fptX*, *pchHI*, and *fepBCDG* reduces the virulence of *Pseudomonas aeruginosa* to *Galleria mellonella* larvae. (A) The bacterial culture of each strain of *P. aeruginosa* shown in the figure was injected into 50 *G. mellonella* larvae. The number of cells injected per *G. mellonella* larva was 10^5^, and data were collected every 12 h. (B, C) *P. aeruginosa* mutant strains (white colonies formed on X-Gal plates) and the PAΔ3Fe strain or Δ*pvdA*Δ*pqsH* strain carrying the *lacZ* gene at the neutral phage attachment site (blue colonies formed on X-Gal plates) were mixed 1:1. The bacterial mixture was injected into the haemocoel of *G. mellonella* larvae, and the hemolymph of *G. mellonella* larvae was collected 24 h later. Competitive index (CI) = colony forming unit (CFU) ratio (white colonies/blue colonies) of the samples after treatment divided by the CFU ratio (white colonies/blue colonies) of the samples before treatment. In-vitro samples were overnight-cultured diluted 1:1000 in Luria–Bertani (LB) medium. All data represent the results of at least three independent experiments. Error bars represent standard deviations. **, *P* < 0.01, ***, *P* < 0.001.

### Interaction network among FptX, PchHI, and FepBCDG

We have demonstrated that FptX, PchHI, and FepBCDG are involved in the uptake of PQS-Fe^3+^ and PCH-Fe^3+^. In addition, FptX can interact with the FepBCDG protein complex. Thus, can PchHI interact with FptX or FepBCDG? Through bacterial two-hybrid assays, this study analyzed the interaction between PchH (or PchI) and FptX, FepB, FepC, FepD, and FepG. The results showed that PchH and PchI had no interaction with FptX, FepB, FepC, FepD, and FepG (Fig. S7). Interestingly, PchH could interact with PchI (Figs. 7A and S7), which was consistent with previously reported results(5). Therefore, it was speculated that PchH and PchI may need to form a heterodimer to interact with FptX, FepB, FepC, FepD, and FepG. As expected, the results of bacterial three-hybrid assays showed that PchHI interacted with FptX, FepB, FepC, FepD, and FepG (Fig. 7B and S7). These results show that there is an interaction between PchHI, FptX, and FepBCDG (Fig. 7C). This suggests that FptX, PchHI, and FepBCDG may form large protein complexes that jointly mediate the uptake of PQS-Fe^3+^ and PCH-Fe^3+^ in *P. aeruginosa*.

**Fig. 7.**
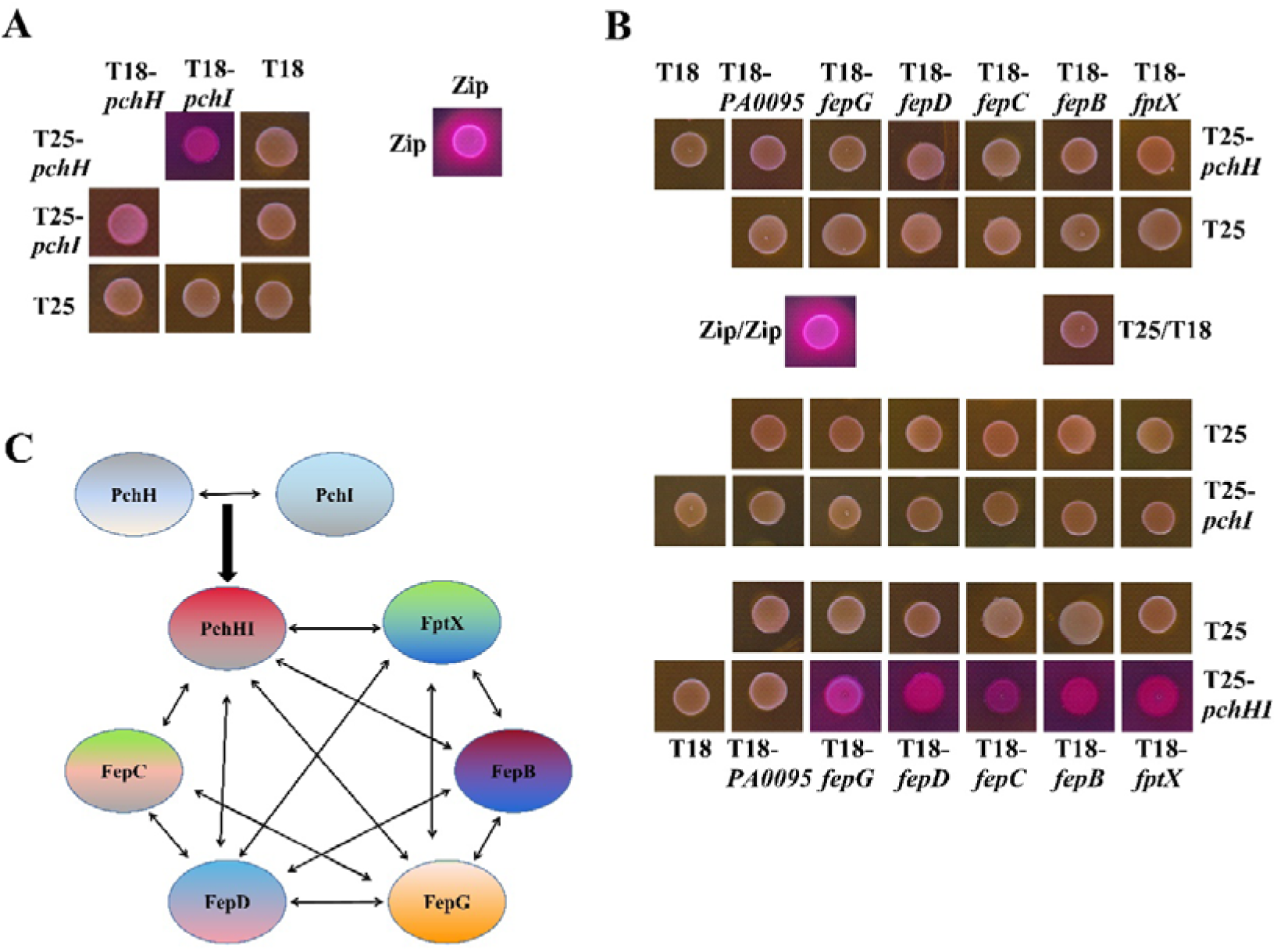
Interactions between FptX, PchHI, and FepBCDG identified by bacterial two-hybrid experiments. (A) PchH interacts with PchI. (B) PchH and PchI form a dimer, PchHI, that interacts with FptX and FepBCDG. (C) The interaction network between inner membrane transporters. Various combinations of recombinant pKT25M and pUT18CM plasmids harboring proteins of interest were co-transformed into *Escherichia coli* BTH101, and the β-galactosidase activity of co-transformants was measured after plating on MacConkey agar plates (Fig. S7). Zip = leucine zipper domain of the yeast transcription factor GCN4 (positive control), T18 = empty vector pUT18CM, T25 = empty vector pKT25M.

## DISCUSSION

*P. aeruginosa* is an opportunistic human pathogen that is listed by the World Health Organization (WHO) as one of the pathogens for which the development of new antimicrobial treatments is urgently needed(51, 52). During infection, *P. aeruginosa* faces stressful environments and must overcome the host immune reactions. In order to survive in these stressful conditions, *P. aeruginosa* secretes a large number of virulence factors, including siderophores(53). Siderophores are small organic compounds produced and secreted by bacteria to uptake iron(13), which is an essential nutrient for bacterial growth and virulence. In the iron uptake system of *P. aeruginosa*, there are three main types of molecules that can be used as siderophores to capture Fe^3+^, namely, PVD, PCH, and PQS. Previous studies have shown that the PQS-and PCH-mediated iron uptake pathways share the same outer membrane transporter, FptA(4, 14), while FptX and PchHI are the inner membrane transporters involved in PCH-mediated iron uptake(5, 14). The present study demonstrates that FptX and PchHI are not only the inner membrane transporters of iron uptake mediated by PCH but also the inner membrane transporters of the PQS-mediated iron uptake pathway. In addition, this study demonstrates that another inner membrane transporter, FepBCDG, also plays an important role in the uptake of PQS-Fe^3+^ and PCH-Fe^3+^ in *P. aeruginosa*.

During the uptake of PCH-Fe^3+^ in *P. aeruginosa*, FptX transports a portion of the PCH-Fe^3+^ into the cytoplasm. FptX is a proton motive-dependent permease that can use proton motive force momentum as its energy source(43). PchH and PchI proteins contain an ATP-binding domain and a transmembrane domain on the same polypeptide(5). This feature corresponds to the YbtPQ and IrtAB ABC transporters of *Yersinia pestis* and *Mycobacterium tuberculosis*, respectively, and both YbtPQ and IrtAB play important roles in the process of iron uptake(54, 55). FepC contains an ATP-binding domain, and this domain provides energy for FepBCDG(7). In the present study, different methods were used to deprive these proteins of their energy source. The results showed that the loss of the energy sources of FptX, PchHI, and FepBCDG led to the inability of *P. aeruginosa* to utilize PQS-Fe^3+^ and PCH-Fe^3+^ (Fig. 4). Interestingly, it was found that the energy of the PchHI transporter complex was derived from the ATPase activity of PchH, but not PchI (Fig. 4D and E).

Promoter-*lacZ* transcriptional fusion assay showed that, unlike FptX, both FepBCDG and PchHI played no role in the autoregulatory loop involving PchR, but further deletion of *fepBCDG* and *pchHI* could reverse the inactive PchR phenotype caused by *fptX* deletion and reactivate the expression of the genes of the PCH pathway under iron-limited conditions (Fig. 5B and D). It has previously been shown that PchR-mediated transcriptional activation of the PCH genes does not require interaction with PCH-Fe^3+^ under very strong iron-limited conditions(56). The deletion of *pchHI* and *fepBCDG* resulted in a similar phenotype, in which PchR also became active in the absence of the obvious transport of PCH-Fe^3+^ into the bacteria. Recent studies have reported that during the uptake of PCH-Fe^3+^ in *P. aeruginosa*, a fraction of the PCH-Fe^3+^ complexes is transported across the inner membrane into the cytoplasm by FptX to interact with PchR in the auto-regulatory loop, while another fraction of the PCH-Fe^3+^ complexes undergo dissociation in the bacterial periplasm via an unknown mechanism, and the free iron is transported further across the inner membrane into the bacterial cytoplasm by PchHI(5). Because FepBCDG and PchHI exhibit similar patterns of effect on PqsR-mediated autoregulatory loop, it is implied that FepBCDG, like PchHI, may act as the ABC transporter to translocate siderophore-free iron into the cytoplasm.

The present study further improves the understanding of the molecular mechanisms of PCH-mediated iron uptake systems. However, there is little knowledge about the molecular mechanisms through which iron is released from the PCH-Fe^3+^ complexes in *P. aeruginosa* cells. It has been reported that FadD1, the fatty acid coenzyme-A ligase, is an interacting partner of the inner membrane transporter FptX, implying that it may play a role in modifying PCH(57). Therefore, we speculate that FadD1 may be involved in the release of iron from the PCH-Fe^3+^ complexes.

The results showed that in TSB medium, the exogenous addition of PQS significantly induced the expression of *phzA1* and *lecA* in *P. aeruginosa* strains PAΔ3FeΔ*pqsA* and PAΔ3FeΔ*pqsA*Δ*fptA*Δ*oprF*Δ*tseF* (Fig. S1). However, in iron-limited medium (TSB medium containing 300 μM 2,2’-bipyridine), the exogenous addition of PQS only weakly activated the expression of *lecA* in *P. aeruginosa* strain PAΔ3FeΔ*pqsA* (Fig. 5A), indicating that PQS may not be efficiently uptaken under iron-limited conditions. Surprisingly, this result was opposite to the results reported by Diggle et al.(38). This different regulatory phenotype may be the result of different culture conditions. An iron-limited CAA medium was used by Diggle et al., and the addition of PQS to this medium could significantly inhibit the growth of *P. aeruginosa* (see Fig. 6 in this article). Additionally, in the present study it was found that whether in iron-rich or iron-limited media, the exogenous addition of PQS-Fe^3+^ could only activate the expression of *phzA1* and/or *lecA* in *P. aeruginosa* strain PAΔ3FeΔ*pqsA*, while it could not activate the expression of these two genes in strains PAΔ3FeΔ*pqsA*Δ*fptA*Δ*oprF*Δ*tseF* and PAΔ3FeΔ*pqsA*Δ*fptX*Δ*pchHI*Δ*fepBCDG* (Figs. S1 and 5A). These results suggest that the function of PQS-Fe^3+^-mediated quorum-sensing regulation is dependent on the TseF-FptA/OprF pathway and three inner membrane transporters, namely FptX, PchHI, and FepBCDG.

This study also investigated whether special organizational structures formed between FptX, PchHI, and FepBCDG. Previous studies have shown that PchH and PchI form a heterodimer, PchHI, which further forms a multiprotein complex with FptX to participate in the uptake of PCH-Fe^3+^(5). However, unlike previous reports that T25-PchH (but not T25-PchI) interacted with FptX-T18(5), neither T18/T25- PchH nor T18/T25-PchI could not interact with T25/T18-FptX and T25/T18- FepB/C/D/G, respectively(Fig. S7). This indicates that the interaction region between PchH and FptX is located at the C-terminus of PchH and the N-terminus of FptX. Interestingly, the present study found that PchHI, a heterodimer formed by PchH and PchI, could interact with FptX and the FepBCDG complex consisting of FepB, FepC, FepD, and FepG (Figs. 7A, 7B, and S7). In addition, FptX also interacted with the FepBCDG complex (Figs. 1D and S2). These results suggest that FptX, PchHI, and FepBCDG may form a larger multiprotein complex that participates in the uptake of PQS-Fe^3+^ and PCH-Fe^3+^ in *P. aeruginosa*.

Based on the results in this study, we proposed a model for *P. aeruginosa* to transport PQS-Fe^3+^ and PCH-Fe^3+^ across the inner membrane into the cytoplasm (Fig. 8). This model also prompted us to put forward the following new hypothesis: although the secretion pathway of PQS and PCH is still unknown, they may share the same secretion pathway, and they may function synergistically. Further work will be required to verify this hypothesis.

**Fig. 8.**
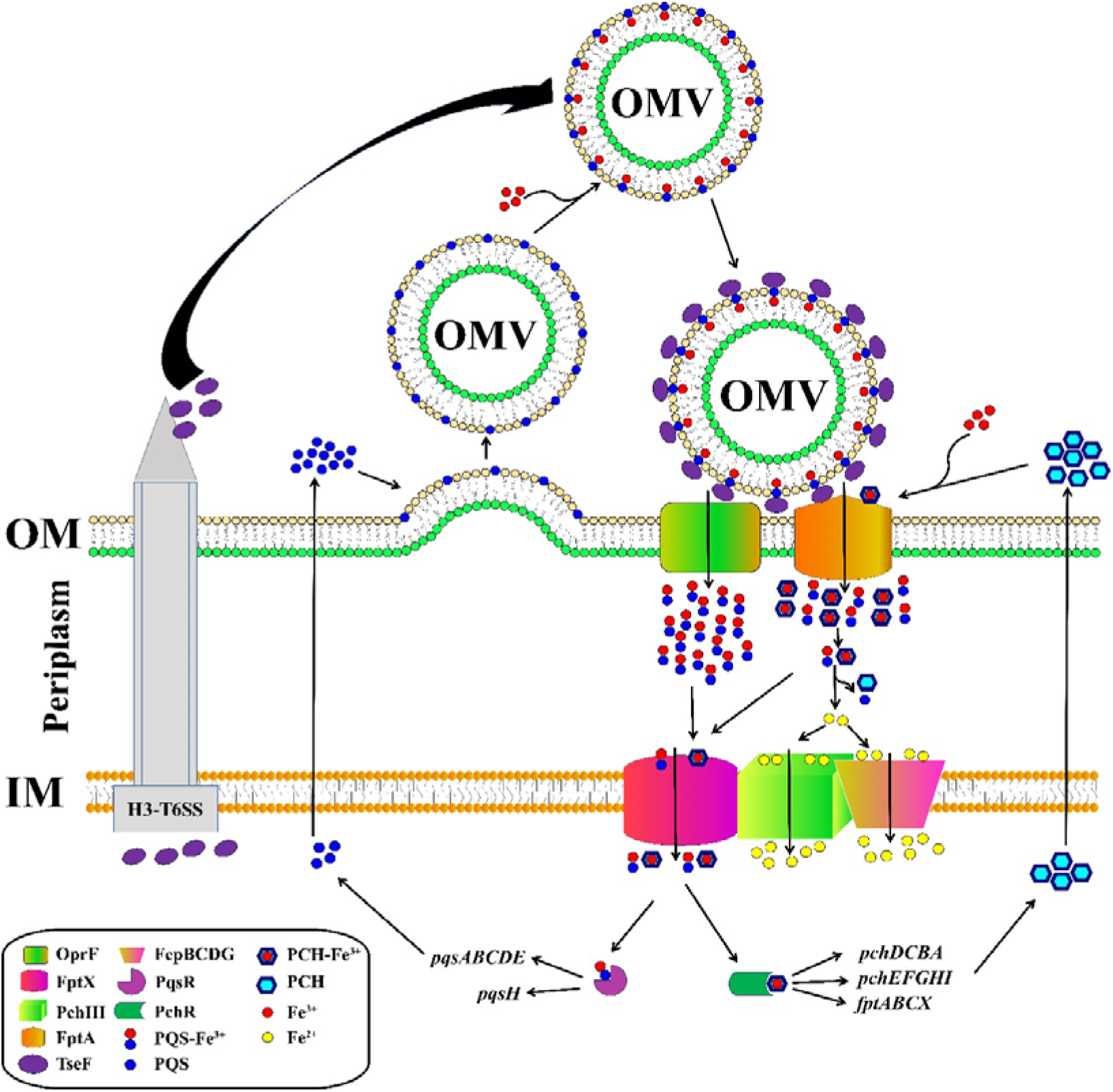
Schematic diagram of the uptake of PQS-Fe^3+^ and PCH-Fe^3+^ by *Pseudomonas aeruginosa*. Extracellular PQS-Fe^3+^ is transported through outer membrane vesicles (OMVs). Under the mediation of the type-VI secretion system (T6SS) effector protein TseF, PQS-Fe^3+^ on OMV enters the periplasm through the outer membrane receptors OprF and FptA. Similarly, extracellular PCH-Fe^3+^ also enters the periplasm through FptA. Then, a fraction of the PQS-Fe^3+^ and PCH-Fe^3+^ complexes enter the cytoplasm directly through FptX, a fraction of the PQS-Fe^3+^ and PCH-Fe^3+^ complexes undergoes dissociation in the bacterial periplasm via an unknown mechanism, and the free iron is transported further across the inner membrane into the cytoplasm by PchHI and FepBCDG. The entry of PQS-Fe^3+^ and PCH-Fe^3+^ into the cytoplasm activates the expression of the PQS and PCH genes to facilitate their synthesis.

Previous studies have shown that *pchHI* is an important virulence gene in *P. aeruginosa*. When *pchHI* mutated, the virulence of *P. aeruginosa* toward *Dictyostelium*, *Drosophila*, and mice significantly decreased(58), indicating that PchH and PchI are necessary for the virulence of *P. aeruginosa*(59). The results of the present study are consistent with these previous reports. Here, it was found that when *fptX*, *pchHI*, and *fepBCDG* were deleted, the virulence of *P. aeruginosa* toward *G. mellonella* larvae significantly decreased. However, the virulence of *P. aeruginosa* was restored to varying degrees after complementing these gene (Figs. 6 and S6), suggesting that FptX, PchHI, and FepBCDG are not only involved in the uptake of PQS-Fe^3+^ and PCH-Fe^3+^ but also play crucial roles in the virulence of *P. aeruginosa*.

In conclusion, PQS and pyochelin in *P. aeruginosa* share inner membrane transporters, including FptX, PchHI, and FepBCDG, to mediate iron uptake. These findings provide a special perspective for the prevention and treatment of *P. aeruginosa* infection, and greatly expand the current understanding of bacterial adaptation to complex environments.

## MATERIALS AND METHODS

### Bacterial strains and growth conditions

The bacterial strains and plasmids used in this study are listed in Supplementary Table S1. The *Escherichia coli* strains were grown at 37°C in either LB or TSB medium. The *P. aeruginosa* strains were grown at 37°C in either LB, TSB, or succinate MM(60). The *P. aeruginosa* PAO1 strain was the parent strain of all of the derivatives used in this study. To construct in-frame deletion mutants, the pK18*mobsacB* derivatives were transformed into relevant *P. aeruginosa* strains through *E. coli* S17-1-mediated conjugation and were screened as described by Lin et al.(4, 61). For overexpression or complementation in the various *P. aeruginosa* strains, the pME6032 derivatives were transformed into the relevant *P. aeruginosa* strains and induced by the addition of 1 mM isopropyl-β-D-1-thiogalactopyranoside (IPTG). Antibiotics were used at the following concentrations for *P. aeruginosa*: kanamycin (30 μg/mL), chloramphenicol (30 μg/mL), gentamicin (200 μg/mL), and tetracycline (160 μg/mL for liquid growth or 200 μg/mL for solid growth). Antibiotics were used at the following concentrations for *E. coli*: kanamycin (30 μg/mL), gentamicin (10 μg/mL), ampicillin (100 μg/mL), chloramphenicol (30 μg/mL), and tetracycline (20 μg/mL).

### Plasmid construction

The construction of the knock-out plasmid was modified from a previously reported study(4, 61, 62). Briefly, to construct the recombinant suicide plasmids for deletion, for the *fptX* gene, the 803-bp upstream and 783-bp downstream fragments flanking the *fptX* gene were amplified with the primer pairs *fptX* Up F/*fptX* Up R and *fptX* Low F/*fptX* Low R, respectively (Supplementary Table S2). The upstream and downstream polymerase chain reaction (PCR) fragments were ligated using overlapping PCR, and the resulting PCR products were inserted into the XbaI/HindIII sites of the suicide vector pK18*mobsacB* to yield the plasmid p-*fptX*. The gentamicin resistance cassette from p34s-Gm was subsequently inserted into the same HindIII site of p-*fptX* to yield the recombinant suicide plasmid pK18-Δ*fptX*. The recombinant suicide plasmids pK18-Δ*pchHI*, pK18-Δ*fepBCDG*, and pK18-Δ*fepC* were constructed in a similar manner using primers listed in Supplementary Table S2.

To construct the complementation plasmid pME6032-*fptX*, PCR-amplified *fptX* was inserted into the EcoRI/BglII sites of the pME6032, giving rise to the recombinant plasmid pME6032-*fptX*. The recombinant plasmids pME6032-*pvdA*, pME6032-*pchHI*, pME6032-*fepBCDG*, and pME6032-*fepC* were constructed using the same method.

The *lecA–lacZ* transcriptional fusions were constructed via the PCR amplification of the 1036-bp upstream DNA region from the *lecA* gene using the primer pairs *lecA* F/*lecA* R (Supplementary Table S2). The PCR amplification products were cloned directly into the pMini-CTX*::lacZ* vector (62), yielding *lacZ* reporter constructs. The recombinant plasmids *fptA*-*lacZ* were constructed using the same method (Supplementary Table S1).

To construct the recombinant plasmids pUT18CM-*fptX* and pKT25M-*fptX*, which were used in the bacterial two-hybrid assays, the gene coding for FptX protein was PCR-amplified using the primer pair two-hybrid *fptX* F/two-hybrid *fptX* R (Supplementary Table S2) and the genomic DNA from *P. aeruginosa* PAO1 as a template. Amplified DNA fragments were digested with BamHI/SalI or BamHI/XohI and subcloned into the corresponding sites of the engineered pUT18-derived pUT18CM (BamHI/SalI) and engineered pKT25-derived pKT25M (BamHI/XohI) vectors, yielding the recombinant plasmids pUT18CM–*fptX* and pKT25M–*fptX*, respectively. The recombinant plasmids pUT18CM–*pchH*, pUT18CM–*pchI*, pUT18CM–*fepB*, pUT18CM–*fepC*, pUT18CM–*fepD*, pUT18CM–*fepG*, pUT18CM–*pchHI*, pKT25M–*pchH*, pKT25M–*pchI*, pKT25M–*fepB*, pKT25M–*fepC*, pKT25M– *fepD*, pKT25M–*fepG*, and pKT25M–*pchHI* (Supplementary Table S1) were constructed using the same method. All constructions were verified using DNA sequencing.

### Site-directed mutagenesis

To construct the site-directed mutagenesis complementary vector pME6032- *pchH*I*, the conservative amino acid residue glutamate on the Walker B motif of the ATPase domain of PchH were replaced with alanine. The 1563-bp front section and 2003-bp rear section DNA segments of the *pchHI* gene were amplified using the primer pairs SD-*pchH*I* Up F/SD-*pchH*I* Up R and SD-*pchH*I* Low F/SD-*pchH*I* Low R, respectively, and the upstream and downstream segments were connected to form a gene segment using overlapping PCR. The product (containing the native Shine-Dalgarno (SD) sequence) of overlapping PCR was inserted into the KpnI/BglII sites of the pME6032 plasmid to yield the site-directed mutagenesis recombinant plasmid pME6032-*pchH*I*. Using the same method, this study constructed pME6032- *phHI\** and pME6032-*fepC\** (Supplementary Table S1).

### Growth assay

The growth assay protocol was as described previously with some modifications(17). *P. aeruginosa* strains were grown overnight in TSB, the overnight cultures were harvested, and the cells were washed with MM twice to remove nutrient-rich substances prior to subculture. Subculture proceeded in MM with 2,2’- bipyridine (250 or 750 µM) with or without IPTG (1 mM), PQS-Fe^3+^ (20 µM), FeCl_3_ (6.7 µM), or PCH (20 µM) to a final OD_600_ of ∼0.01. Cultures were incubated at 37°C and OD_600_ readings were taken every 12 h for 72–120 h.

### Construction and screening of the bacterial two-hybrid library

The protocol used to construct genome fragment libraries was as described previously with some modifications(63, 64). The genomic DNA of *P. aeruginosa* strain PAO1 was prepared using the PureLink Genomic DNA kit (Invitrogen, Carlsbad, CA, USA) and partially digested by S*au3A*I. The randomly digested DNA was separated on a 0.8% agarose gel, and fragments ranging in size from 1,000 to 3,000 bp were gel-purified using the Qiagen gel extraction kit (Qiagen, Valencia, CA, USA). The genomic DNA libraries were constructed using the pKT25M vectors. The pKT25M vectors were digested with B*amH*I and dephosphorylated with phosphatase. The pools of DNA fragments were ligated overnight at 16°C into the different pKT25M linearized vectors using T4 ligase (NEB, Ipswich, MA, USA). The resulting ligation mixture was transformed into *E. coli* TG1 competent cells. The libraries were collected and pooled as the prey libraries and stored in a freezer at −80°C.

The plasmid of pUT18CM-*fptX* was used as the bait to probe the genomic DNA libraries. Basically, 25–50 ng of each pKT25M-derived library was transformed into 100 μL of electrocompetent BTH101 cells carrying the pUT18CM-*fptX* bait vector and plated on MacConkey agar medium containing 0.5 mM IPTG. Bacteria expressing interacting hybrid proteins will form red colonies on MacConkey agar medium, while cells expressing non-interaction proteins will remain white. IPTG was used to increase β-galactosidase expression. A co-transformant containing pKT25-*zip* and pUT18-*zip* was used as a positive control for expected growth on the screening medium. A co-transformant containing empty vector pKT25 and pUT18 was used as a negative control. The red colonies were picked up and recultivated in liquid medium, and plasmids were isolated and further analyzed using DNA sequencing.

### Bacterial two-hybrid assay

Bacterial two-hybrid assays were performed using previously described methods(65, 66). In brief, pUT18CM and pKT25M carrying different genes were used in various combinations to co-transform *E. coli* BTH101 cells, and the plate was cultured at 30°C for 24 h. Five independent colonies were selected and inoculated into LB liquid culture medium supplemented with 100 µg/mL ampicillin, 30 µg/mL kanamycin, and 0.5 mM IPTG. After overnight growth at 30°C, 3 µL of each culture was spotted onto MacConkey plates supplemented with 100 µg/mL ampicillin, 30 µg/mL kanamycin, 0.5 mM IPTG, and 1% maltose, then cultured for 20 h at 30°C. The formation of red colonies on MacConkey agar plates indicated an interaction between the two proteins, and white colonies indicated negative results.

### β-galactosidase assay

The β-galactosidase assays were modified from a previously reported study(4, 61). A total of 100 µL of bacterial culture was added to 900 µL of Z Buffer (40 mM NaH_2_PO_4_, 10 mM KCl, 60 mM Na_2_HPO_4_, 1 mM MgSO_4_, and 0.2% β-mercaptoethanol). A total of 1 µL of 0.1% sodium dodecylsulfate (SDS) and 50 µL of chloroform were added to the suspension, which was mixed vigorously for 20 s. The suspension was then incubated for 1 h at 30°C. A total of 100 µL of 4 mg/mL 2- nitrophenyl β-D-galactopyranoside (ONPG) (Sigma, St. Louis, MO, USA) was added to the cells. The reaction was stopped by adding 500 µL of 1 M Na_2_CO_3_. The suspension was centrifuged at 10,000 × g for 3 min, and the absorbance of the supernatant was read at 420 and 550 nm using a microplate reader. The β- galactosidase activity was then calculated in Miller units (MUs) according to the following equation:

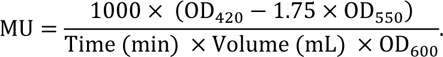

### Extraction of *P. aeruginosa* PCH

The extraction of *P. aeruginosa* PCH was modified from a previously reported study(67). Briefly, *P. aeruginosa* was grown in 1-L volumes of MM medium at 37°C, 200 rpm for 48 h. The bacterial cells were removed by centrifugation (6,000 g for 15 min at 23°C), and the supernatant fluid was brought to pH 1–2 with 1 M HCl. Ethyl acetate was added in a 1:5 ratio, and after vigorous shaking in separatory funnels the ethyl acetate layers were collected and concentrated using rotary evaporation. The residue remaining in the flask after evaporation was dissolved in 1 mL methanol. When PCH extracts were used, it was added to the culture medium at a ratio of 1: 1000.

### In vitro co-culture assay

In vitro co-culture assays were modified from a previously reported study(68). *P. aeruginosa* strains Δ*pvdA*Δ*pqsH*Δ*pchE*Δ*fptX*Δ*pchHI*Δ*fepBCDG* (pBBR1MCS-5), PAΔ3FeΔ*pqsA*Δ*fptX*Δ*pchHI*Δ*fepBCDG* (pBBR1MCS-5), PAΔ3FeΔ*pqsA* (pME6032), and Δ*pvdA*Δ*pqsH*Δ*pchE* (pME6032) were cultured in 5 mL of TSB liquid medium (37°C, 200 rpm, and 20 h). One milliliter samples of different strain cultures with the same OD_600_ were centrifuged at 4°C and 3,000 g for 10 min. As much supernatant was removed as possible, and the bacterial cells were retained. One milliliter of fresh MM medium containing kanamycin was added and the bacterial cells were resuspended. The cells were centrifuged at 4°C, 3,000 g for 10 min to remove as much supernatant as possible, and the bacterial cells were retained. The cells were resuspended in 1 mL of fresh MM medium containing kanamycin, and the OD_600_ values of different cultures were adjusted to the same level in order to start with the same number of cells. PAΔ3FeΔ*pqsA* (pME6032) and Δ*pvdA*Δ*pqsH*Δ*pchE*Δ*fptX*Δ*pchHI*Δ*fepBCDG* (pBBR1MCS-5) were mixed in a 1:1 ratio, as well as Δ*pvdA*Δ*pqsH*Δ*pchE* (pME6032) and Δ*pvdA*Δ*pqsH*Δ*pchE*Δ*fptX*Δ*pchHI*Δ*fepBCDG* (pBBR1MCS-5), to produce a mixed bacterial suspension. The mixed bacterial suspension was diluted at a ratio of 1:1000 in a fresh MM medium containing kanamycin (if needed, appropriate amounts of 2,2’- bipyridine, PQS-Fe^3+^, PCH, or hemin were included in the medium), and culture was conducted at 37°C and 200 rpm. Samples were collected at regular intervals and diluted to 10^−7^ using a 10-fold dilution method. Three microliters of each dilution were collected and spotted onto LB plates supplemented with kanamycin and tetracycline or kanamycin and gentamicin. Incubation was conducted at 37°C for 2 days. The colonies were counted and the number of colony-forming units (CFUs) in 1 mL was calculated. Finally, the growth curve was plotted using log10 (CFU/mL). All assays were performed in triplicate.

### Determination of intracellular metal ion content

The intracellular metal ion content determination method was modified from reference(69, 70). Briefly, *P. aeruginosa* strains were cultured in 5 mL of TSB liquid medium at 37°C and 200 rpm for 20 h. After 1 mL culture solutions were collected and washed twice with MM, cells were subcultured in MM medium at a ratio of 1:100 until the exponential phase, centrifuged at 4°C and 2,000 g for 10 min, and the bacterial cells were collected. PBS buffer containing 1 mM ethylene diamine tetraacetic acid (EDTA) was used to suspend the bacterial cells, suspensions were centrifuged at 2,000 g for 10 min at 4°C, the bacterial cells were collected, and the process was repeated once. The bacterial cells were suspended using PBS buffer, centrifuged at 4°C and 2,000 g for 10 min, and collected. The bacterial cells were resuspended in PBS and divided into four equal parts. Then, 0.4% glucose and 20 µM PQS-Fe^3+^ (20 µM PCH or 20 µM hemin) were added to each part. Each portion was incubated at 37°C and 200 rpm for 0, 1, 2, and 4 h, and centrifuged at 4°C and 2,000 g for 20 min to collect the bacterial cells. PBS buffer containing 1 mM EDTA was used to suspend the bacterial cells, suspensions were centrifuged at 4°C and 2,000 g for 20 min, the bacterial cells were collected, and the process was repeated once. The cells were washed again with PBS buffer, centrifuged at 4°C and 2,000 g for 20 min, and collected. The wet cell pellet weight was determined and bacteria were chemically lysed using 5 mL Bugbuster (Novagen, Madison, WI, USA) (gram wet pellet cell paste)^−1^ according to the manufacturer’s instructions. Bacterial cells were resuspended in Bugbuster solution by pipetting and incubation on a rotating mixer at a slow setting for 20 min. The total protein content of each sample was measured using a BioRad protein assay (BioRad, Hercules, CA, USA) according to the manufacturer’s instructions. The wet pellet weight and total protein content for each sample were noted. Each sample was diluted 100-fold in 3% molecular-grade nitric acid to a total volume of 10 mL. Samples were analyzed using inductively coupled plasma mass spectrometry (ICP-MS) (Varian 802-MS; Varian, Palo Alto, CA, USA), and the results were corrected using the appropriate buffers for reference and dilution factors. Triplicate cultures of each strain were analyzed during a single experiment and the experiment was repeated at least three times.

### *G. mellonella*-killing assay

The protocol for the *G. mellonella-*killing assay was as described previously with some modifications(71). *P. aeruginosa* strains were cultured in 5 mL of TSB liquid medium at 37°C and 200 rpm overnight. Subculture was conducted in 5 mL of fresh TSB containing kanamycin at a 1:100 ratio until the OD_600_ value reached 0.5. Bacterial cells were collected via centrifugation at 4°C and 3,000 g for 5 min. The bacterial cells were suspended in 0.85% NaCl solution, centrifuged at 4°C and 3,000 g for 5 min, and collected, and the procedure was repeated twice. The bacterial cells were suspended and diluted with 0.85% NaCl solution to a cell count of 2 × 10^7^ CFU/mL. The *G. mellonella* larvae were placed on ice for 5 min to put them under anesthesia. A micro syringe was used to inject 10^5^ cells into the haemocoel of 3-day-old, fifth-instar *G. mellonella* larvae, and 0.85% NaCl solution was injected as a control. In each group, 50 *G. mellonella* larvae were injected and cultured at 25°C in the dark, and the procedure was repeated with three groups for each strain. Data were recorded every 12 h. Data were analyzed using Kaplan–Meier survival curves. Statistical significance was assessed using the Mantel–Cox log rank test, applying Bonferroni’s correction for multiple comparisons.

### *G. mellonella* co-infection experiments

A *lacZ* reporter gene was transferred to the neutral phage attachment site (*attB*) of the *P. aeruginosa* chromosome as follows: the recombinant plasmid pMini-CTX-P*tac*::*lacZ*(4) was transformed into *E. coli* S17-1, and the resulting plasmid was then transferred to the *P. aeruginosa* chromosome (*attB* site) by mating and selection for tetracycline resistance. The selectable marker was removed by the transient expression of the Flp recombinase from plasmid pFLP2, which was then cured by counter-selection on sucrose plates (72). The resulting *P. aeruginosa* strain Δ*pvdA*Δ*pqsH attB*∷P*tac*-*lacZ* and existing PAΔ3Fe *attB*::P*tac*-*lacZ*(4) were confirmed to grow as well as Δ*pvdA*Δ*pqsH* and PAΔ3Fe in both in-vitro and in-vivo competition assays.

The *G. mellonella* co-infection experiment was performed as previously described with minor modifications(4, 73). *P. aeruginosa* strains were cultured in 5 mL of TSB liquid medium at 37°C and 200 rpm for 20 h. Cells were subcultured in 5 mL of fresh TSB medium at a ratio of 1% until the exponential phase, centrifuged at 4°C and 3,000 g for 10 min, and collected. The bacterial cells were suspended in PBS buffer, centrifuged at 4°C and 3,000 g for 10 min, and collected. This step was repeated twice. The bacteria were suspended and diluted with PBS buffer to a cell count of 2×10^7^ CFU/mL. In each experiment, a *P. aeruginosa* strain that contained *lacZ* reporter inserted at the neutral phage attachment site (producing blue colonies on X-Gal plates; 40 μg/mL) was mixed 1:1 with a *P. aeruginosa* strain without *lacZ* (producing white colonies on X-Gal plates). For *G. mellonella* co-infection, the *G. mellonella* larvae were placed on ice for 5 min to put them under anesthesia. Five microliters of a mixed bacterial suspension with a CFU of 1×10^5^ were injected into the haemocoel of 3-day-old, fifth-instar *G. mellonella* larvae, and 0.85% NaCl solution was injected as a control. After 24 h, the hemolymph of eight infected larvae from each group was collected in 1.5-mL Eppendorf tubes containing 2 μL of 1% phenylthiourea (PTU) on ice. The samples were serially diluted with sterile PBS and spread on LB agar plates containing kanamycin and 40 μg/mL X-Gal. Culture was conducted overnight at 37°C and cultures were left at 22–25°C until blue colonies appeared. In vitro co-culture: the mixed bacterial suspension was subcultured in fresh LB liquid medium containing kanamycin at a ratio of 1:1000 at 37°C and 200 rpm for 12 h. The samples were serially diluted with sterile PBS and spread on LB agar plates containing kanamycin and 40 μg/mL X-Gal. Culture was conducted overnight at 37°C and cultures were left at room temperature until blue colonies appeared. Both the total CFU and the ratio of blue-to-white bacteria were determined. For the competitive index (CI) calculation: CI = CFU ratio of the sample after treatment (white colonies/blue colonies) / CFU ratio of the sample before treatment (white colonies/blue colonies). The larvae were selected randomly for each test group.

### Statistical analysis

All of the experiments were performed in triplicate and repeated on two different occasions. The data are expressed as the mean ± S.D. The differences between the frequencies were assessed using Student’s *t*-test (bilateral and unpaired), and a *p*-value of 0.05 was considered to be statistically significant. The Shapiro–Wilk test and one-way analysis of variance were performed using the GraphPad Prism version 7.00 software (GraphPad software Inc.; San Diego, CA, USA) to examine the normality of the data and the homogeneity of the variances, respectively. GraphPad Prism 7 and Adobe Illustrator 2020 (CS6; Adobe, Mountain View, CA, USA) were used to create all of the figures.

### Data availability

The authors declare that all the relevant data supporting the findings of this study are available within the article and its Supplementary Information files or from the corresponding author on request.

## ACKNOWLEDGMENTS

This work was supported by the National Natural Science Foundation of China (31700031, 32070103), the Qinchuang Yuan “Scientist + Engineer” Team Construction Project of Shaanxi Province (2023KXJ-019), the Regional Development Talent Project of the “Special Support Plan” of Shaanxi Province 2020-44, a grant from the Outstanding Young Talent Support Plan of the Higher Education Institutions of Shaanxi Province 2018-111, and the Youth Innovation Team of Shaanxi Universities 2022-943.

